# The coordination of replication initiation with growth rate in *Escherichia coli*

**DOI:** 10.1101/2021.10.11.463968

**Authors:** Anna Knöppel, Oscar Broström, Konrad Gras, David Fange, Johan Elf

**Affiliations:** Department of Cell and Molecular Biology, Science for Life Laboratory, Uppsala University, Uppsala, Sweden

## Abstract

*Escherichia coli* coordinates replication and division cycles by initiating replication at approximately the same size per chromosome at all growth rates. By tracking replisomes in individual cells through thousands of division cycles, we have dissected the mechanism behind this precise process. We have characterized wild-type cells grown under different conditions and also many mutants related to the expression and binding states of the initiator protein DnaA. This rich data set allowed us to compare the relative importance of all previously described control systems. We found that the replication initiation size regulation is not strongly dependent on the absolute concentration of DnaA, nor does it depend on active *dnaA* expression. Replication initiation is also not consistently triggered by cell division or replication termination. In contrast, some of the factors that convert DnaA between its ATP- and ADP-bound states have a strong effect on initiation size. We suggest a plausible model for DnaA-ATP mediated triggering of initiation at fast growth, where regulatory inactivation of DnaA (RIDA) is the main system for monitoring the number of chromosomes during active replication.

## Introduction

All cells need to coordinate DNA replication with cell growth such that each chromosome is, on average, replicated once per generation. This is particularly challenging for bacteria with a wide range of generation times, some of which are even shorter than the time it takes to replicate the chromosome. The ambition of this work is to understand how *E. coli* adjusts replication initiation control to its growth rate. This is a classical field of study which has been approached from different directions over the decades.

Through the mechanistic genetic/biochemical path, a number of mechanisms that contribute to regulation have been described, including *e*.*g*. the critical role of DnaA-ATP for initiating replication at *oriC* (Hansen and Atlung, 2018), autoregulation of *dnaA* expression (Hansen and Rasmussen, 1977), SeqA-dependent sequestration of newly replicated *oriC* to prevent re-initiation (Lu et al., 1994), and binding of DnaA-ATP to DnaA boxes throughout the chromosome (Schaper and Messer, 1995). A number of systems have also been described to modulate DnaA’s state of ATP or ADP binding; the chromosomal locus *datA* and the replication machinery associated Hda protein both reduce the initiation potential by supporting the hydrolysis of DnaA-ATP to DnaA-ADP (Kasho and Katayama, 2013; Katayama et al., 1998; Kato and Katayama, 2001; Kitagawa et al., 1996; Kurokawa et al., 1999). Conversely, the chromosomal loci *DARS1* and *DARS2* (Fujimitsu and Katayama, 2004; Fujimitsu et al., 2009) convert DnaA-ADP to the apo-form. The apo-form then quickly binds ATP, as there is more ATP than ADP in the cell (Bochner and Ames, 1982). From here on out, we will write it as if the conversion goes straight from DnaA-ADP to DnaA-ATP. Membrane-associated acidic phospholipids have also been described to serve the same function as the *DARS* sites (Sekimizu and Kornberg, 1988). It is, however, still not known how these systems operate together and if they are sufficient to regulate replication initiation, *i*.*e*. to trigger initiation when there is too little DNA compared to cell mass.

There is also a more recent phenomenological path that has emerged from the breakthroughs in live-bacterial imaging, initiated by the invention of the mother machine microfluidic chip (Wang et al., 2010). This path was initially focused on statistical properties of cell size variation from generation to generation since this was what could be measured with phase-contrast microscopy alone (Amir, 2014; Campos et al., 2014; Taheri-Araghi et al., 2015). It is, however, clear that these studies could only define limits that mechanistic models should satisfy; it is important to complement the imaging studies with molecular reporters, for example, by tracking replication forks with fluorescence microscopy (Si et al., 2019; Wallden et al., 2016), which has previously only been done with low-throughput methods (Bates and Kleckner, 2005; Lau et al., 2003; Li et al., 2002).

In this study, we have combined the two paths and studied replication and division cycles using high-throughput fluorescence microscopy while at the same time perturbing the system to gain insights into the importance of each component. We began by characterizing the coordination of replication to the division cycle at different growth rates, then assessed the importance of *dnaA* regulation and expression level, and finally examined the relative contributions of the various systems modulating the DnaA-ATP and -ADP bound states. The results are evaluated in relation to previously described models for control of replication initiation (Box 1).

### Box 1: Possible mechanisms for balancing replication initiation to cell growth

**Box 1 Figure.**
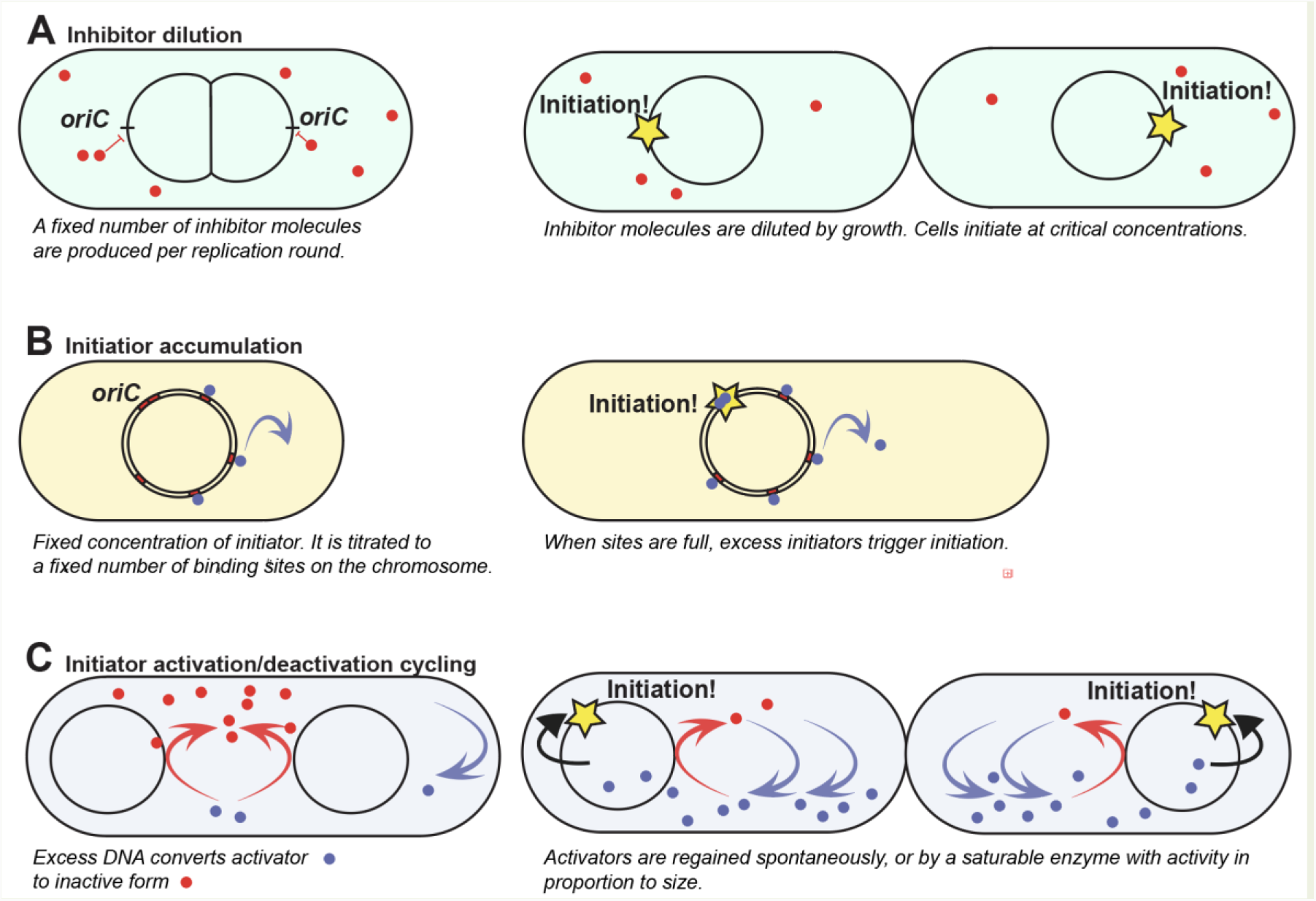
Plausible models for coordination between growth and replication initiation. (**A**) Inhibitor dilution model. (**B**) Initiator accumulation model. (**C**) Initiator activation /deactivation cycling model.

On average, cells trigger a new round of replication on each origin once per generation. The cell size at replication initiation has little variation (Sauls et al., 2019), and independently of growth rate and cell size, the ratio of chromosomal DNA to protein stays within 25% (Dennis and Bremer, 1974). At least four principally different mechanisms that are capable of achieving this level of regulation have been proposed, all of which are based on triggering replication when there is too little DNA compared to the cell size.

#### A. Inhibitor dilution

In this hypothetical mechanism, each copy of the genome expresses a constant number of replication inhibitors per unit of time, *i*.*e*. expression of the replication inhibitors is limited by transcription and not translation and the concentration of RNA polymerase is constant. The inhibitors are diluted by growth such that initiation is triggered when the repressor concentration drops below a specific value (D’Ario et al., 2021; Schmoller et al., 2015; Zatulovskiy et al., 2020). This mechanism relies on a very sensitive concentration dependence for the repression and would be exceptionally sensitive to noise in gene expression. A similar model was proposed by (Pritchard et al., 1969) and suggests that for each replication event, a fixed number of inhibitors are made, which are then diluted by growth.

#### B. Initiator accumulation

For these types of models, the initiator is kept at a constant concentration in the cell such that the number of molecules is proportional to the cell size. This can be achieved, for example, by expressing the initiator as a constant fraction of the proteome instead of in proportion to the concentration of chromosomes, or by utilizing autoregulated gene expression (Hansen et al., 1991; Sompayrac and Maaloe, 1973). The initiators bind strongly to a fixed number of sites per chromosome (Hansen et al., 1991; Roth and Messer, 1998). When these sites are full, excess synthesis contributes to increasing the concentration of free initiators, and thus, initiation is triggered at a fixed volume per chromosome each generation. This model could, in principle, be implemented by the binding of DnaA to chromosomal DnaA boxes.

#### C. Initiator activation/deactivation cycling

In this class of models, the initiator is cycled between an active and an inactive state, where the flux to the active state increases with increasing volume and the flux to the inactive state increases with the number of chromosomes. If the modification reactions are saturable, triggering at a fixed chromosome to volume ratio can be made ultra-sensitive (Koshland et al., 1982; Wallden et al., 2015).

#### D. Mechanical models

There is also a class of initiation models where replication is triggered based on other processes in the cell cycle, effectively moving the regulatory problem to these processes. For example, if replication is triggered after cell division, the challenge is not to initiate at a set size, but to divide at a set size. It is also possible to envision replication being triggered by replication termination,which moves the regulatory problem to replicating the chromosome at a rate that is proportional to the growth rate.

## Results

### Experimental setup and data visualization

Our typical experimental procedure for studying replication in relation to the cell cycle is to grow *E. coli* MG1655 cells in a mother machine-type fluidic chip that allows for rapid medium swaps (Baltekin et al., 2017). We use this strategy to monitor thousands of individual cell cycles in each experiment. Growth is imaged by phase-contrast and replication forks are localized by detecting the fluorescence (Figure 1A) of YFP translational fusions to DnaN, DnaQ, or SeqA. DnaN and DnaQ are both parts of the replisome (O’Donnell, 2006), and SeqA binds hemimethylated GATC in the wake of the complex (Lu et al., 1994). The different replication markers gave very similar information (Figure S1A). The localization of replication forks is presented in so-called fork plots, which are either shown with multiple generations or one generation (Camsund et al., 2020). The fork plots show where along the cell’s long axis the replication forks are located at a particular cell size. We refer to the densest regions of the fork plots as branches, where the start of each branch is interpreted as the initiation of individual rounds of DNA replication with two replication forks moving bidirectionally away from the origin region (Wallden et al., 2016). For the fork plots with multiple generations, size is written as if division events were disregarded. We refer to this type of structure as a “super-cell”. Termination of replication is interpreted as being when each branch ends abruptly. In most of the fork plots, the average cell sizes at birth and division are indicated with dashed white lines. The methods section describes how these quantities were defined and calculated, but importantly, the individual replication forks were tracked in time (Figure 1A) such that, replication initiation, for example, could be automatically assigned in most cases based on the start of the tracks of detected foci linked together by u-track (Jaqaman et al., 2008). Cells were grown at 30 **°**C in M9 minimal medium supplemented with a carbon source and 1×RPMI amino acids solution unless otherwise stated. The growth medium is referred to only by the name of the carbon source; precise media compositions can be found in Table S1.

**Figure 1.**
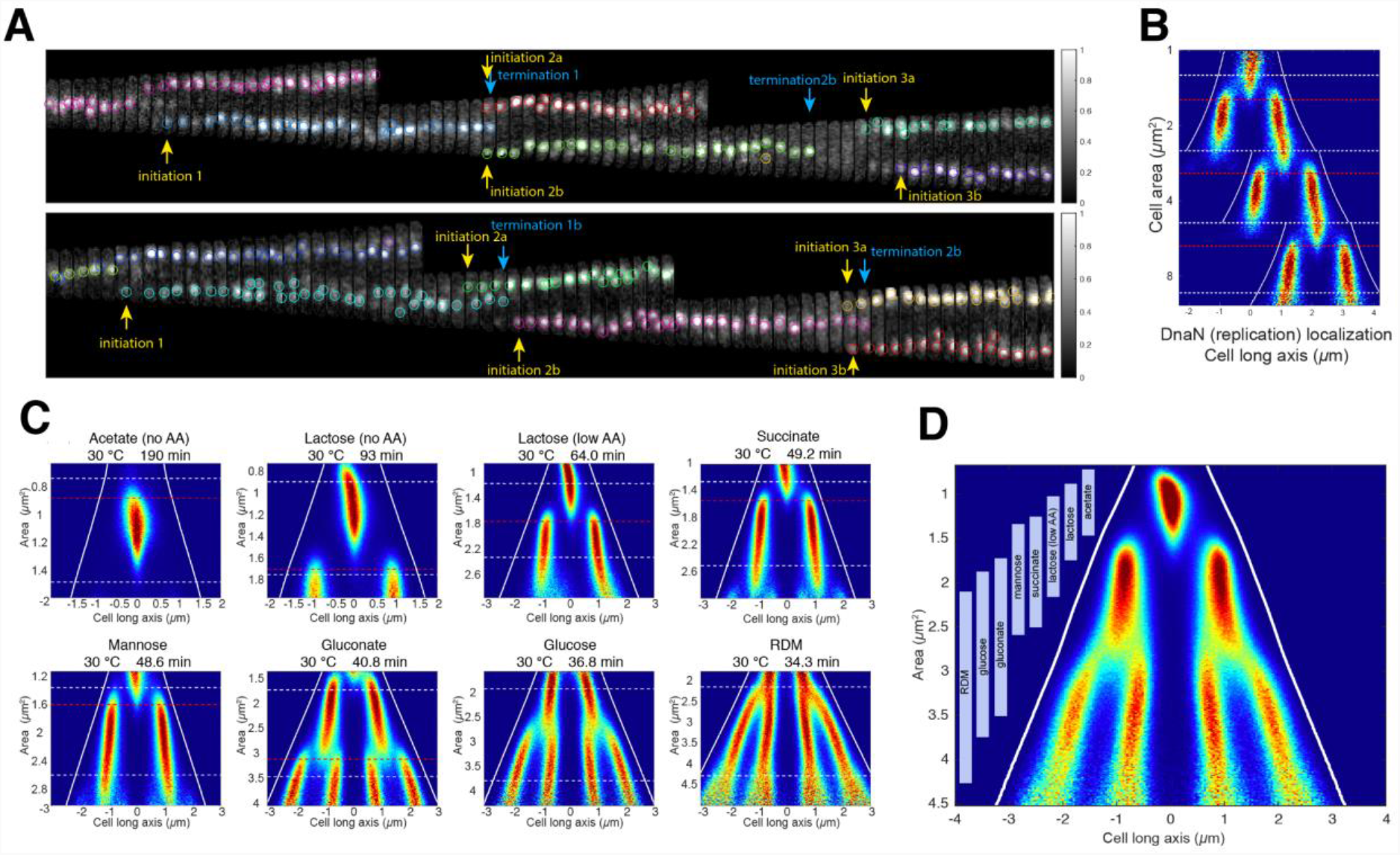
Replication patterns in different growth media. (**A**) Examples of cells tracked over three generations. YPet-DnaN foci are marked with circles where different colors indicate one concurrent replication, i.e. two replication forks that often are spatially co-localized. Cell outlines are displayed with a gray line. (**B**) Three generation fork plot. A two-dimensional distribution of position of YPet-DnaN foci along the long axis of the cells (x-axis) and cell size (y-axis) is plotted. Here, the cell size in generations two and three are defined as if the cells in the previous generation(s) had not divided. Dashed white horizontal lines indicate the average sizes of cells at birth or division. Dashed horizontal red lines indicate the median initiation size as determined by tracking replisomes in single cells. Solid white lines indicate average cell pole positions. (**C**) Fork plots for strains carrying YPet-DnaN grown in different media. Medium, growth temperature, and corresponding doubling time is given above each fork plot. Lines as in (B), but here red dashed lines indicate the average instead of the median. (**D**) Pooled fork plot with the data from (C) including the same number of cells from each growth condition. Bars indicate the division cycle of each medium.

### Spatial organization of replication is not dependent on growth condition or cell division

To quantify the effect of the growth medium on replisome organization, we imaged YPet-DnaN-expressing cells in eight different growth media (Figure 1C). We found that the cells appeared to use the same guiding principle to spatially organize their replication regardless of the growth medium, although it differed in the number of concurrent replication processes. On average, cells grown in acetate without amino acid supplementation initiated replication on one origin, cells grown in lactose (without and with 2.5% supplementation of amino acids), succinate, and mannose concurrently initiated on two origins, and cells grown in gluconate, glucose, and RDM initiated on four. To highlight the similarities in replication organization, we pooled the data from all different growth media and constructed a fork plot for the entire set in a common coordinate system (Figure 1D). Apart from the cells grown in RDM, the majority of the cells initiated their replication at a similar ratio of initiation size to the number of concurrently initiated origins, giving the pooled fork plot a well-groomed appearance (Figure 1D).

Since initiation can occur right before division (*e*.*g*. for cells grown in gluconate), or right after division (*e*.*g*. for cells grown in mannose), division appears unimportant for triggering replication at intermediate growth rates (Table S2). This is also in line with a recent report from the Jun lab (Si et al., 2019).

### Initiation of DNA replication can be triggered in the absence of both transcriptional regulation and expression of the *dnaA* gene

When we tracked replication forks at the level of single cells, we observed correlation coefficients of 0.3–0.5 for replication initiation to cell size, independently of growth conditions (Table S2). This is in line with previous observations (Si et al., 2019). A correlation coefficient of 0.5 is typical of an adder phenotype where an, on average, constant and independent cell size increment is added between two subsequent initiations. The adder phenotype can arise if the cell accumulates an initiator molecule proportionally to its volume and initiates replication when it has accumulated a fixed number of initiator molecules (Ho and Amir, 2015; Si et al., 2019). This corresponds to the initiator accumulation model in Box1. The natural candidate for the initiator is DnaA; DnaA bound to ATP can initiate DNA replication *in vitro* (Sekimizu et al., 1987) and the intracellular concentration of DnaA has been shown to negatively correlate with cell size at initiation (Atlung and Hansen, 1993; Si et al., 2019). Additionally, DnaA-ATP is known to autoregulate its own transcription (Hansen and Rasmussen, 1977).

To investigate whether autoregulation of DnaA transcription is important for triggering initiation at a fixed volume, we constructed a strain in which the expression of a sole copy of *dnaA* is constitutive. This was done by introducing *dnaA* under the control of a constitutive promoter (J23106) at a neutral position and exchanging the two native promoters of the *dnaA* operon as well as the *dnaA* gene with a constitutive promoter (J23106) in front of the remainder of the operon (*dnaN, recF*, and *gyrB*). The *P*_*J23106*_*-dnaA* strain showed identical average expression of DnaA as compared to wild-type (wt; Figure S2) and a minor increase in initiation and division sizes as well as in variation of replication initiation volumes (Figures 2A and S2). It displayed an indistinguishable growth rate compared to an isogenic reference strain, both when grown in the microfluidic chip and in a plate reader (Figures S2A and S2D). We concluded that the possibility of DnaA to control its own expression does not have a critical role in the regulation of replication initiation. However, this result does not rule out the gene expression-based model for initiator accumulation since *dnaA* gene expression in proportion to cell size could potentially be achieved by other means (Box 1).

**Figure 2.**
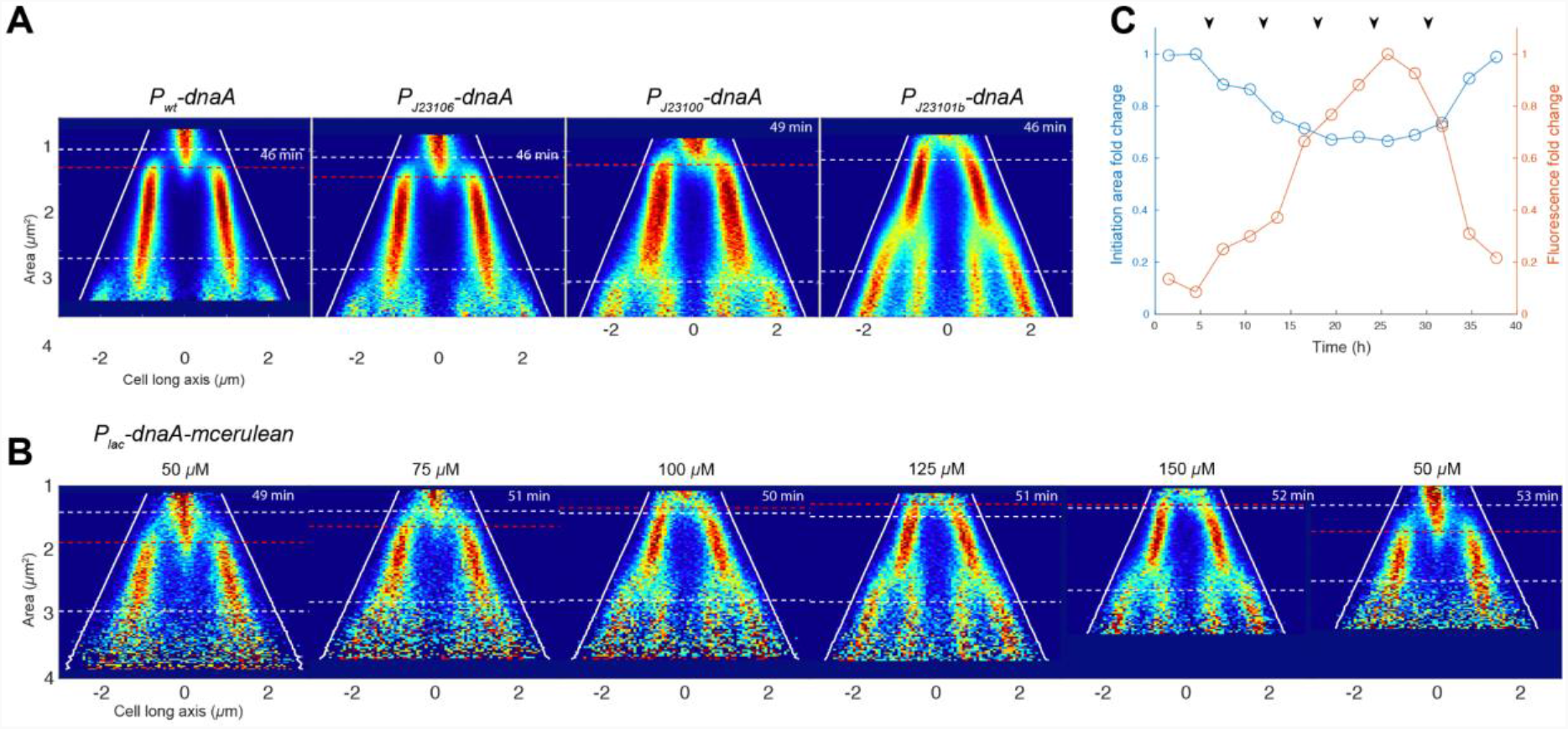
Replication initiation at different DnaA concentrations. (**A**) Fork plots for SeqA-Venus in strains carrying the native promoter of dnaA (P_wt_-dnaA), a constitutive promoter with the same DnaA expression level (P_J23106_-dnaA), and two strains with higher expression levels as compared to wt (P_J23100_-dnaA and P_J23101b_-dnaA). The expression levels of DnaA are found in Figure S2A. Lines as in Figure 1C. (**B**) Fork plots for a SeqA-Venus in strain carrying the IPTG controllable P_lac_ promoter in front of dnaA transcriptionally fused to mCerulean (P_lac_-dnaA-mCerulean) from an experiment where the IPTG concentration in the growth medium was changed stepwise. IPTG concentrations are given above each fork plot. The cells were grown in succinate medium at 30 **°**C. Lines as in Figure 1C. (**C**) Fold change of mCerulean fluorescence and initiation size over time for the experiment shown in (B). Black arrows indicate when the IPTG concentration was changed.

The non-critical effect of removing DnaA’s transcriptional autoregulation did, however, allow us to modify the DnaA expression level by placing *dnaA* behind other constitutive promoters or the isopropyl β-D-1-thiogalactopyranoside (IPTG) controllable *lac* promoter. When expressing *dnaA* through the *lac* promoter, we transcriptionally fused the cyan fluorescent protein mCerulean to allow for measurements of DnaA expression. In a version of the initiator accumulation model where gene expression supplies the initiator molecule, we expect increased gene expression to result in smaller initiation sizes. This model states a 1:1 relationship between gene expression rate and initiation size; a two-fold increase in gene expression rate means that the threshold is reached faster and the new initiation size is halved compared to before the increase. In line with the initiator accumulation model and previous observations (Atlung and Hansen, 1993; Si et al., 2019), we found that the initiation size divided by the number of origins on which initiation occurred correlated negatively with DnaA expression, both for the IPTG-inducible strain (Figures 2B and 2C) and for the strains with different constitutive promoters (Figures 2A and S2A). Furthermore, the increased DnaA expression had no significant effect on growth rate or cell size (Figures 2 and S2), implying an increased DNA concentration when DnaA was overexpressed. However, in conflict with the gene expression-based initiator accumulation model, we found that the five-fold increase in DnaA concentration mediated by the *P*_*lac*_ at high IPTG concentrations only resulted in a 35% decrease in initiation size.

As a more definitive test of the role of gene expression in the initiator accumulation model, we investigated if new synthesis of DnaA is needed for replication initiation. In an initiator accumulation model based on gene expression, shutting down gene expression would prohibit further initiations. To test this prediction, we used the strain described above with a single copy of *dnaA* under the control of the *lac* promoter and transcriptionally fused to the fluorescent protein mCerulean. RT-qPCR measurements of the *dnaA* transcript before and after removing IPTG showed that *dnaA* was effectively turned off when IPTG was removed (Figures S3A and S3B). In the IPTG titration above, we found that too low expression of DnaA led to growth arrest (Figure S3C). Under the microscope, we observed replication initiation arrest and that the cells became filamentous as their growth rate decreased (Movie S1). To distinguish whether cells stopped initiating replication due to lack of expression or due to low intracellular concentration of DnaA, we supplied DnaA in excess before turning off its gene expression (Figure 3). In this way, we still had time to observe possible initiations before the DnaA concentration became too low. To corroborate that *dnaA* expression was turned off, we measured the mCerulean cell fluorescence signal per area after removal of IPTG. We found that the time required to dilute mCerulean two-fold (52 min) corresponded to the generation time in the replication experiment (49 min), suggesting effective repression of DnaA expression. We note that our proteomic LC-MS/MS analysis gave a diverging result for protein dilution (Table S6), but we consider the mCerulean measurements more accurate since they were acquired in the same experiment as the initiation size measurements.

**Figure 3.**
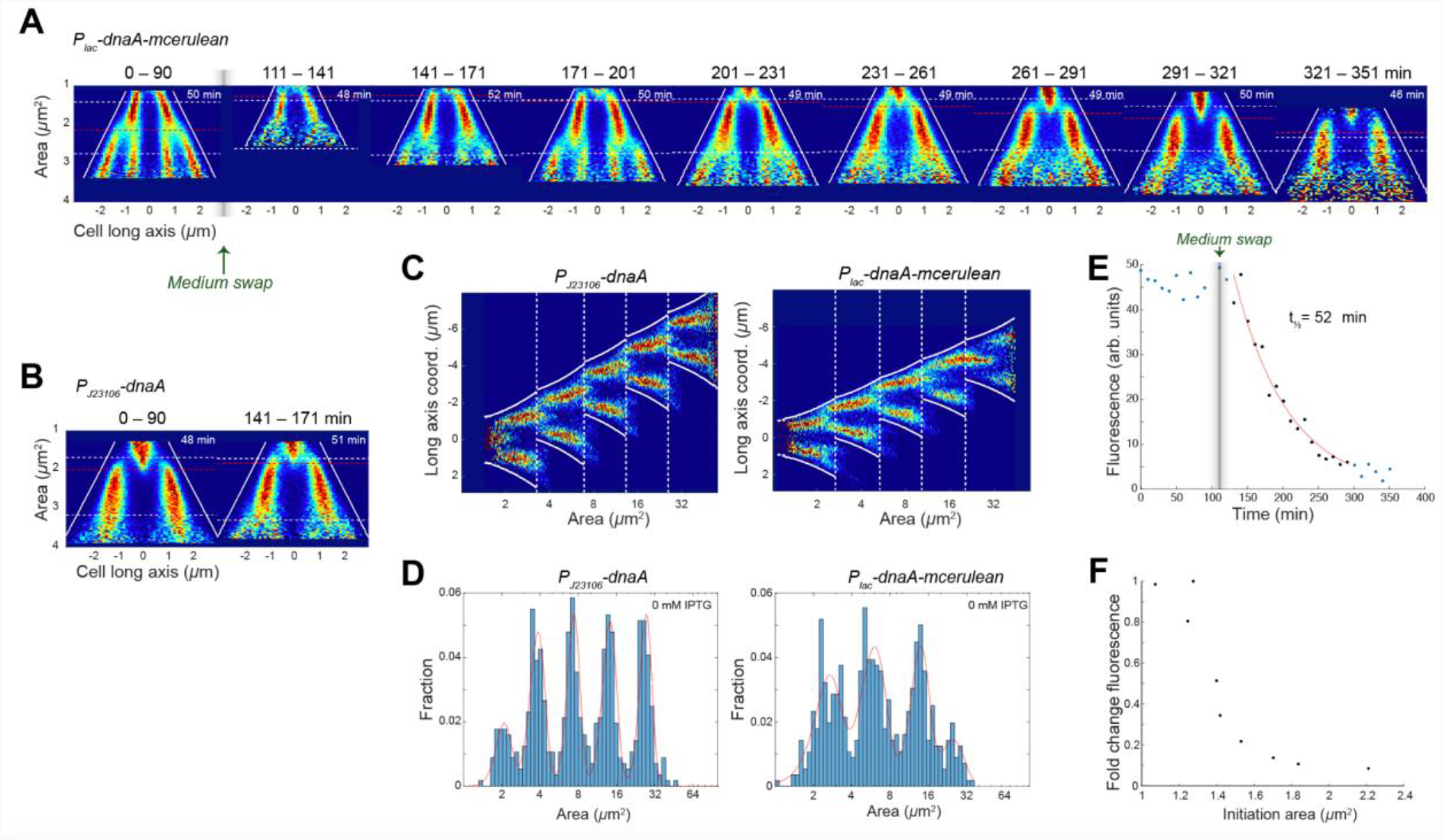
Replication initiation after turning off expression of dnaA. **(A)** Fork plots for a strain with IPTG controllable DnaA expression (P_lac_-dnaA-mCerulean) before (0–90 min) and after (111–351 min) the medium was swapped from medium with 1 mM IPTG to medium without IPTG (see detailed procedure in the methods section). Lines and numbers as in Figure 1C. **(B)** As in (A) but for a strain with constitutive expression of DnaA (P_J23106_-dnaA) grown in the same microfluidic chip. One representative fork plot after the medium swap (141–171 min) is shown. **(C)** Five generation fork plot after medium swap (111–351 min) for constitutive (left) and IPTG regulated (right) dnaA expression. **(D)** Distribution of single-cell initiation areas (blue bars) of the same cells as in (C). Cells were a part of the same lineage and located at the bottom of the channels. Red lines show the result of a regression of the distribution to either a sum of five (left) or a sum of four (right) Gaussians. **(E)** Mean of the per cell mCerulean fluorescence intensity area density as a function of time both before and after the medium swap. Background fluorescence from cells not expressing mCerulean were subtracted from each cell intensity density (see method sections for details). At each time point, a new set of cells were imaged to avoid the effect of photobleaching. Time points after the medium swap (black) were fitted to a single exponential decay (red). **(F)** Initiation area over fold change fluorescence after medium swap.

After turning off DnaA synthesis, the cells gradually initiated replication at larger and larger size-to-chromosome ratios as the DnaA concentration decreased by dilution. ∼200 minutes, or ∼4 generations, after DnaA synthesis was turned off, the initiation size in the IPTG inducible mutant was the same as wt. Within the next generation following this point, the low DnaA concentration could not sustain regular initiations. (Figure 3; repeat experiment Figures S3D–F). For further verification, we tracked cells at the bottom of the microfluidic channels for five generations and saw that they were able to repeatedly initiate replication in the same manner as the whole population shown in Figures 3C and 3D. In line with the weak dependence of intracellular DnaA concentration on initiation size during the stepwise increase of DnaA expression rate (Figures 2 and S2), we found that a 50% reduction in DnaA concentration due to dilution by growth only resulted in a 15% increase in initiation size (Figure 3F). We concluded that the result from turning off DnaA expression contradicts gene expression-based models where accumulation of DnaA to a threshold level triggers initiation. This would, for example, rule out the specific model where new synthesis of DnaA is titrated to binding sites that have stronger affinity than *oriC*, allowing initiation to occur only when these sites are filled (*i. e*. the initiator accumulation model; Hansen et al., 1991).

Furthermore, the fact that the chromosome concentration was relatively constant when *dnaA* was significantly overexpressed implies that excess DnaA over chromosomal binding sites alone does not trigger initiation. The presence of separated peaks of initiation sizes in consecutive generations (Figure 3D) after DnaA expression was turned off instead indicates that cells did not initiate replication directly after origin sequestration had ended, *i*.*e*. 15 min after initiation (Campbell and Kleckner, 1990). This implies that excess DnaA is somehow prevented from initiating replication after origin sequestration has ended, possibly due to the presence of an inhibitor or because DnaA has been converted into a non-active form. DnaA-ADP could have been a plausible inhibitor of initiation, but, although DnaA-ADP cannot initiate replication by itself, it has been shown to promote rather than inhibit initiation when present together with DnaA-ATP (Crooke et al., 1992; Leonard et al., 2019; Yung et al., 1990). RIDA, on the other hand, is a well-documented example of a system that converts the active initiator form into the inactive (Katayama et al., 1998), even in strains with high overexpression of DnaA. Direct measurements of DnaA-ADP and DnaA-ATP concentrations in bulk samples also suggest that DnaA-ADP dominates (Kurokawa et al., 1999). It seems most likely that the reason excess DnaA does not trigger initiation is that most of the molecules are inactive.

### Deletion of the *hda* gene gives a growth rate-dependent initiation phenotype

The Hda protein has been shown to be necessary for RIDA functionality (Kato and Katayama, 2001). To investigate the consequences of RIDA inactivation, we constructed a Δ*hda*::*kanR* mutant in a SeqA-Venus labeled strain. The *hda* deletion-strain grew indistinguishably from wt in succinate medium, but very poorly when the medium was swapped to LB (Figure 4B). We used whole genome sequencing to ensure that the *hda* deletion strain did not contain any of the previously described compensatory mutations (Charbon et al., 2011). While the division sizes in the *hda* deletion mutant were similar to wt (Figure 4A), the initiation size to chromosome ratio was substantially smaller than wt, with the fork plot appearing similar to the case where DnaA was overexpressed (Figures 2B and 3A). When the medium was swapped from succinate to either RDM or LB, the initiation size became undefined (Figures 4A, S4E and S4F) and the cell growth rate decreased gradually until growth stopped (Figure 4B). A growth defect is also what we would expect if Hda was the main mechanism to convert DnaA to ADP-form, since a *hda* deletion would result in significant over-initiation. This is corroborated by our results (Figures 4A, 4B, S4E and S4F), since the growth rate and initiation accuracy decreased significantly in LB and RDM compared to the less nutritious succinate medium. That deletion of *hda* results in over-initiation, is also in line with previous reports that overexpression of the origin sequestration protein SeqA can suppress loss of Hda (Charbon et al., 2011).

**Figure 4.**
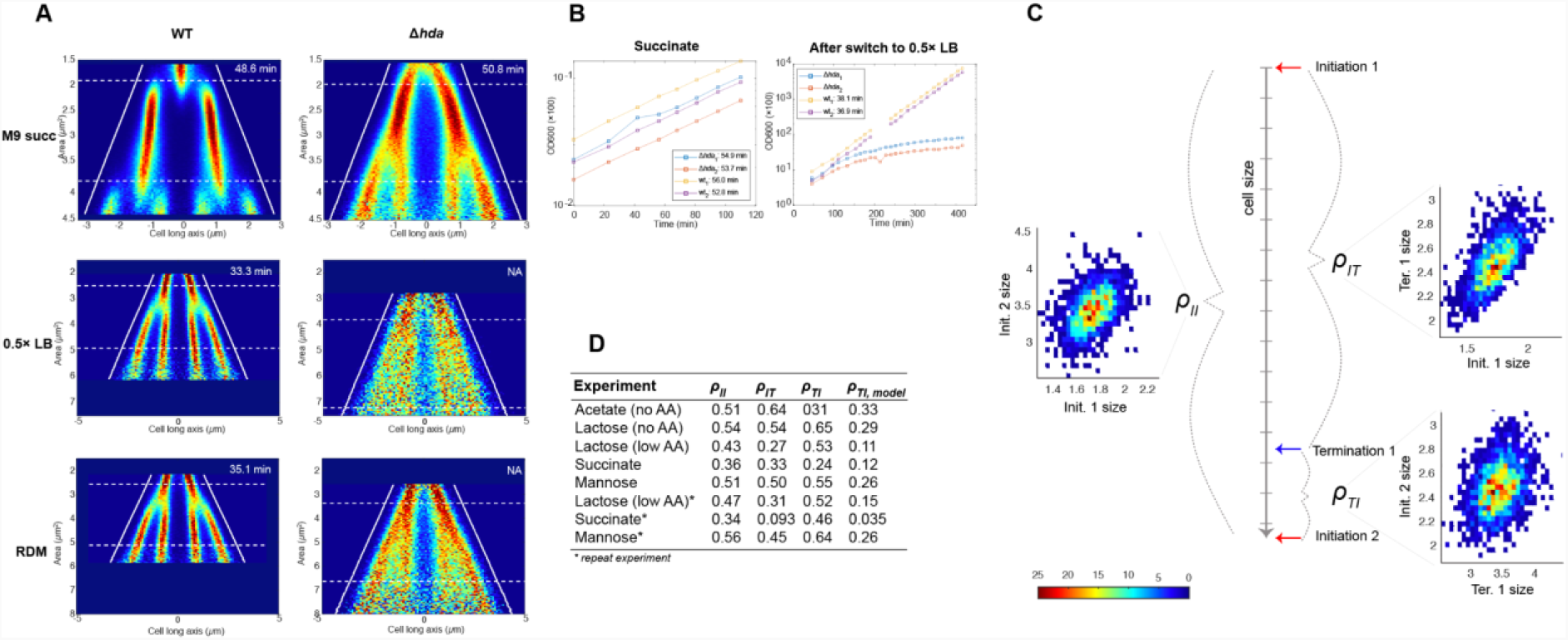
Replication phenotype of Δhda and cell size correlation coefficients between initiation and termination events. **(A)** Fork plots of Δhda and wt strains grown in succinate, 0.5× LB, and RDM media. For the cases of 0.5× LB and RDM the cells were initially kept in succinate medium and then swapped to either RDM or 0.5× LB. The fork plots shown are cells imaged between 160–240 min after the medium swap. The full experiments are presented in Figures S4E and S4F. **(B)** Batch growth of Δhda and a wt strain grown first in succinate medium then swapped to 0.5× LB. **(C)** Cartoon of the time span between two initiations. The cell size correlation between two consecutive initiation events (ρ_II_), initiation and termination of the same round of replication (ρ_IT_), and the termination of one replication round and initiation of the next coming round (ρ_TI_) are depicted and showcased in the figure with the YPet-DnaN strain grown in acetate medium (Figure S4B for more conditions). **(D)** Cell size correlation coefficients as defined in the main text from the YPet-DnaN strain grown in various media.

RIDA is dependent on active replication (Katayama et al., 1998). This implies that if there are competing processes converting between the ATP- and ADP-forms of DnaA, the net flux of DnaA-ATP production should increase following termination of replication and possibly trigger new initiations. To test if there is such an effect, we correlated termination events to the next initiation event in individual cells (Figures 4C, 4D and S4B). To ensure that we could determine a meaningful termination-to-initiation size correlation, we first made sure that the ends of the time-lapse fluorescent foci trajectories were truly indicating termination events by showing that replisomes and termini (*ter*-mCherry; Figure S3C for fork plots with the different carbon sources) are close in space when the YPet-DnaN trajectories end (Figure S4D).

Interpreting the correlation between termination and initiation is, however, non-trivial, since the connection between correlation and causality is extra complicated for cyclical processes. For example, for cells grown in acetate, we found that sizes at the beginning and end of each replication round, *i*.*e*. from initiation to the corresponding termination, were correlated with a correlation coefficient *ρ*_IT_ ≈ 0.64. We also found the size at initiation and the size at the subsequent initiation were correlated with a correlation coefficient *ρ*_II_ ≈ 0.5 (Figure 4D and Table S2). By modeling the two correlated size pairs as bivariate Gaussian distributions with shared initiation sizes, it is possible to, without adding any additional information, predict the residual correlation, *ρ*_*TI, model*_ between the end of one replication round and the beginning of the next, *i*.*e*. from termination to the next initiation. Using the measured *ρ*_IT_ and *ρ*_II_ for the acetate case, we predicted a correlation coefficient between termination and initiation of *ρ*_*TI, model*_ ≈ 0.33. This is similar to the experimentally measured, where *ρ*_*TI*_ ≈ 0.31, implying that a causal mechanism is not required to explain the observed correlation. For cells grown in richer media where termination and initiation occur closer in time (Figure 1C), the correlations between the end of one replication round and the beginning of the next cannot be explained by the model above (Figure 4D). Instead, the increase in correlation between termination and initiation could potentially be explained by cessation of RIDA at termination.

### Deletion of *DARS* can be compensated for by deleting *data* and overexpressing *dnaA*

In addition to RIDA, we have characterized the importance of other regulatory pathways dependent on DnaA-ATP or DnaA-ADP by mutating the components involved in the DnaA-ATP/ADP conversion. *datA*-dependent DnaA-ATP hydrolysis (DDAH) has been described to be important for DnaA-ATP to ADP conversion (Kasho and Katayama, 2013) and *DARS1, DARS2*, and acidic phospholipids are all suggested to be involved in the conversion of DnaA-ADP into DnaA-ATP (Fingland et al., 2012; Fujimitsu and Katayama, 2004; Fujimitsu et al., 2009; Sekimizu and Kornberg, 1988; Xia and Dowhan, 1995).

To remove the complexity introduced by the autoregulation of DnaA expression, we made deletions of the *DARS1, DARS2*, and *datA* loci, as well as combinations of double- and triple-knockouts, in the strain with constitutive expression of *dnaA* from *P*_*J23106*_*-dnaA*. As expected (Frimodt-Møller et al., 2016; Kasho et al., 2017), the size at which the cells initiated replication was reduced when *datA* was deleted and increased when *DARS1* or *DARS2* were deleted (Figures 5A and S5). The *DARS1* deletion mutant gave rise to a particularly broad distribution of initiation sizes. However, the widening of the initiation size distribution did not cause a growth rate reduction (Figures S5B and S5C), as would have been expected if the large variability in initiation size increased the risk of dividing without finishing replication. That the deletion of *DARS1* gave a stronger phenotypic response as compared to *DARS2* is the opposite of what has been reported previously (Fujimitsu et al., 2009; Kasho et al., 2014). We do not know the reason for this, but we do show that this difference did not arise from the exchange of the wt promoter for DnaA expression to a constitutive one (Figures S5A–C). Importantly, we found that deletion of *datA* (Figures 5A and S5A) could partially revert the *DARS* phenotypes. This suggests that the *DARS1* deletion mutant simply has too little DnaA-ATP and contradicts the hypothesis that the *DARS* and *datA* sites are critical for the replication-initiation control as such. The scenario in the *DARS1* deletion mutant is also clearly different from that with too low DnaA concentration (Figure 3A); the latter produces filamentous cells and dies, whereas the first initiates irregularly but survives.

**Figure 5.**
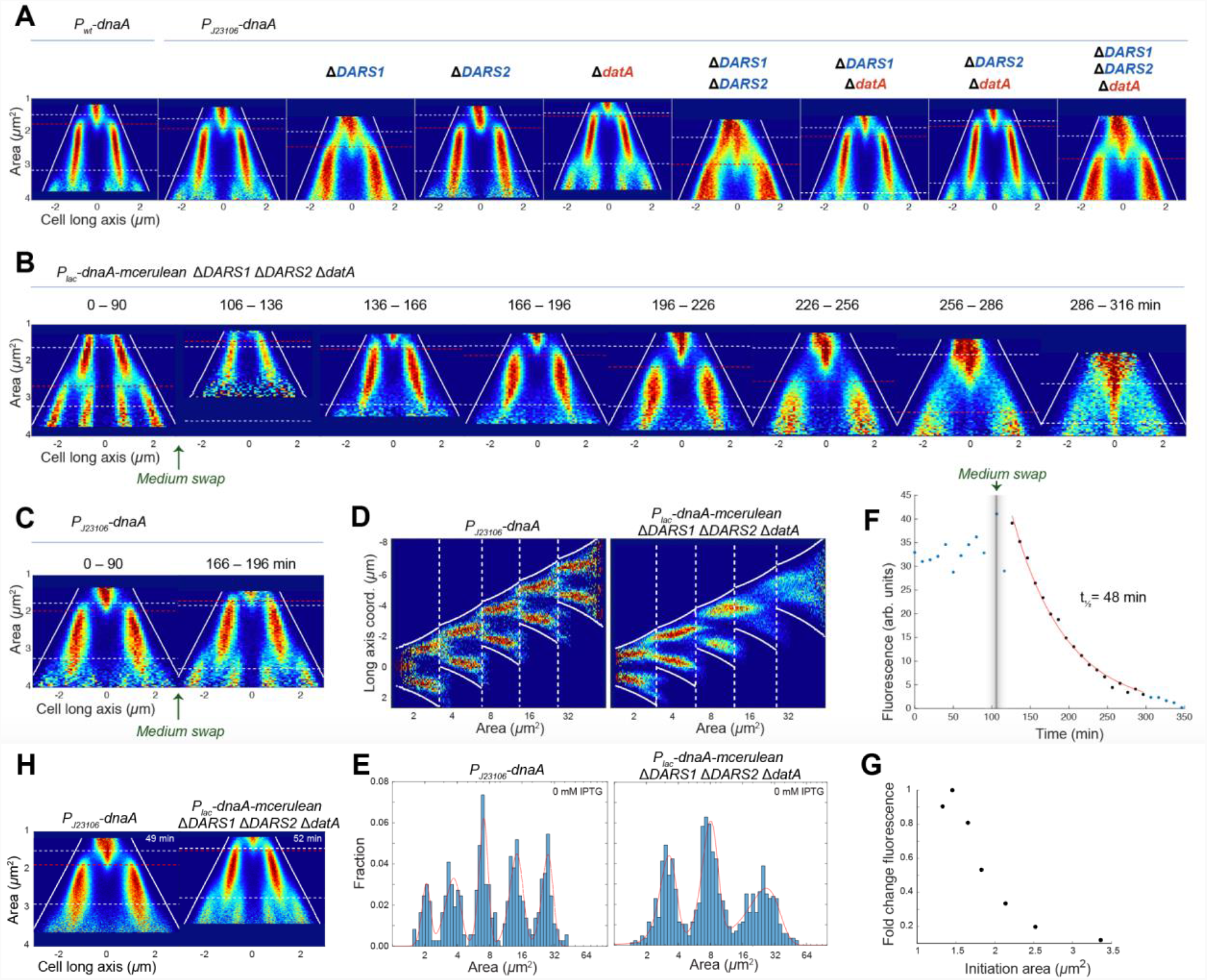
Replication phenotypes of DnaA-ATP/ADP regulatory mutants. (**A**) Fork plots of DnaA-ATP/ADP regulation mutants. The mutations were constructed in the J23106-dnaA background, for similar constructions in dnaA wild-type background see Figure S5A. Blue text indicates ADP- to ATP- form converter and red text indicates ATP- to ADP-form converter. (**B**–**G**) Similar to Figure 3 but for a ΔDARS1 ΔDARS2 ΔdatA mutant with IPTG controllable expression of dnaA and mCerulean (P_lac_-dnaA-mCerulean). (**H**) Expression of dnaA using 100 µM IPTG in a ΔDARS1 ΔDARS2 ΔdatA mutant compared to a reference strain with constitutively expressed dnaA.

As a more definitive test of the importance of *DARS* sites in triggering initiation, we performed the same experiment as above where the expression of *dnaA* was turned off (Figure 3), but this time in a mutant lacking *DARS1, DARS2*, and *datA*. Using an IPTG-inducible *dnaA* construct in the triple (*DARS1:DARS2:datA*)-knockout background, we found that there were multiple new rounds of replication initiated after synthesis of DnaA was stopped, also in the absence of *DARS* and *datA* sites (Figures 5B–G). In addition, we also found that IPTG-induced expression of DnaA can revert the *DARS* deletion phenotypes (Figure 5H), similarly as was observed for the deletion of *datA* (Figure 5A). We conclude that another mechanism than *DARS* must be responsible for converting DnaA-ADP into DnaA-ATP, potentially phospholipid-assisted conversion from the ADP- to the ATP-form.

### Knockdown of *pgsA* does not change the initiation size

Could the source of the DnaA-ATP molecules required to initiate replication be dependent on the acidic phospholipids cardiolipin (CL) and phosphatidylglycerol (PG)? *In vitro* experiments have shown that especially CL can promote dissociation of ADP from the DnaA-ADP complex (Castuma et al., 1993; Sekimizu and Kornberg, 1988). The *pgsA* gene encodes for phosphatidyl glycerophosphate synthase, a protein that catalyzes the biosynthesis of CL and PG (Gopalakrishnan et al., 1986). Temporarily turning off *pgsA* gene expression has been shown to stop growth until *pgsA* expression is turned back on (Fingland et al., 2012; Xia and Dowhan, 1995). Importantly, the effect is not seen in mutants that are not dependent on DnaA for initiation.

We have previously shown that dCas9-mediated repression of *pgsA* expression causes cells to grow ∼30% slower while maintaining the same initiation sizes compared to non-repressed cells (Camsund et al., 2020). The impact on growth rate is admittedly lower than for the *pgsA* deletion mutants (Fingland et al., 2012; Xia and Dowhan, 1995), but if acidic phospholipids are a major contributor to regenerating the ATP-form of DnaA, we would still expect *pgsA* repression to cause increased initiation sizes. Also, since the *DARS* sites are (supposedly) performing the same function as the acidic phospholipids, we would expect *DARS*-deletion mutants to be more sensitive to changes in the acidic phospholipid concentration.

Contradictory, we found that knocking down *pgsA* did not cause any significant change in initiation size in neither a reference strain with constitutively expressed DnaA nor in the *DARS1*:*DARS2* deletion mutant, and the knockdown resulted in a similar growth rate decrease in both strains (Figures 6 and S6). To make sure that we were not selecting for viable cells in the analysis of the microfluidics experiment, we also compared the growth rates in flask cultures (Figures S6B and S6C). Also in these experiments, we found that the reference and the *DARS1*:*DARS2* deletion mutant had the same growth rate reduction as a result of *pgsA* repression.

**Figure 6.**
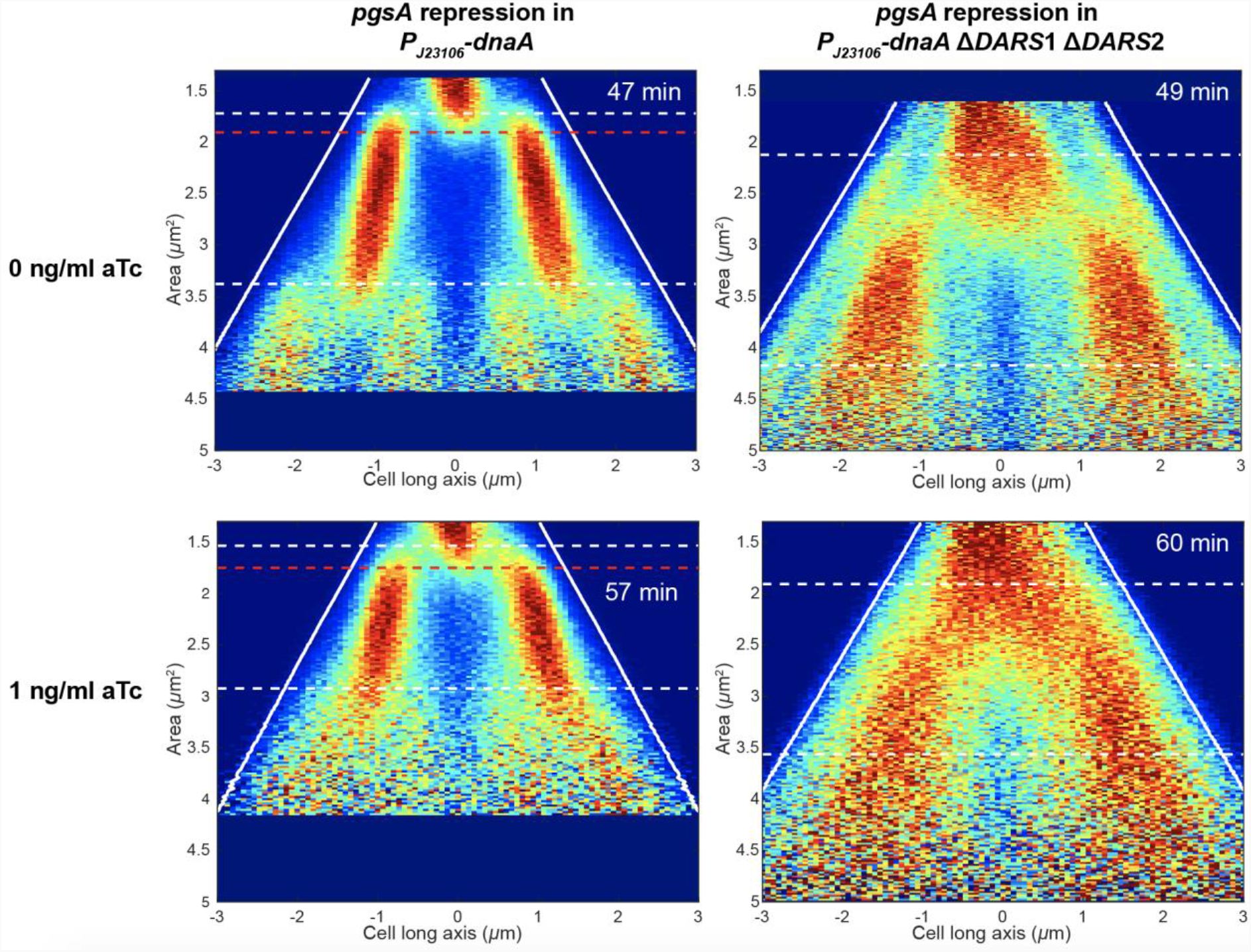
Repression of pgsA. Fork plots of constitutively expressed DnaA and ΔDARS1 ΔDARS2 strains with the dCas9 repression system (Camsund et al., 2020) targeting the expression of pgsA. Two experiments are shown where the two strains were run in parallel: uninduced dCas9 (0 ng/ml aTc) and fully induced (1 ng/ml aTc). More conditions and strains are presented in Figure S6A.

## Discussion

The results of our experiments shed light on the feasibility of the models described in Box 1. First, we can rule out mechanistic models based on triggering initiation just after cell division or replication termination since there are growth conditions where initiation typically occurs just before these events.

Considering the inhibitor dilution models, the only candidate inhibitor we can think of would be DnaA-ADP, which may compete with DnaA-ATP for binding critical sites in *oriC*. DnaA-ADP is a plausible candidate since it can be made in a stoichiometric relation to the number of chromosomes, for example if DnaA-ATP is expelled from the DnaA boxes and converted to the ADP-form by RIDA, for DnaA-ADP to later be diluted by growth to a trigger concentration. If the DnaA-ADP inhibition is sufficiently potent, it may even explain the weak dependence on DnaA overexpression. However, *in vitro* experiments have shown that DnaA-ADP promotes rather than inhibits initiation when present together with DnaA-ATP (Crooke et al., 1992; Leonard et al., 2019; Yung et al., 1990)

We have designed specific tests of the initiator accumulation model. Increasing or decreasing the total DnaA concentration does not result in a corresponding change in the amount of chromosomal DNA, showing that initiation is not triggered at a specific DnaA to chromosome ratio, at least not in the succinate growth condition. We also find that individual lineages of cells can continue to initiate for several consecutive generations after DnaA synthesis has been turned off if the DnaA expression level was elevated before it was turned off. This shows that new synthesis of DnaA is not needed to trigger initiation, which rules out a gene expression-based initiator accumulation model.

Our results give the strongest support for an initiator activation/deactivation model. In line with Donachie and Blakely, (2003), we find that regulation of replication initiation is primarily mediated by DnaA-ATP or the DnaA-ATP to DnaA-ADP ratio. First, the cell-to-cell variation in initiation volume changes when we disturb mechanisms related to the conversion, *i*.*e. DARS1, DARS2, datA*, and Hda. The *DARS* and *datA* mutations are even compensating for each other, clearly suggesting that it is the state of DnaA-ATP/ADP binding that is important for initiation and not the absolute concentration of DnaA.

At all growth rates, initiation is triggered at a relatively constant chromosome to cell size ratio in successive generations, even at growth rates with multiple, sometimes overlapping replication cycles. A robust implementation to achieve this type of regulation by the ATP/ADP state of DnaA is to use an integrating feedback control (Yi et al., 2000) based on initiator activation/deactivation, *i*.*e*. a modification-demodification cycle (Koshland et al., 1982) where the rate of converting DnaA into its ATP form is proportional to the cell size and the rate of conversion into the ADP-form is proportional to the number of chromosomes. A similar feedback is implemented by using the aminoacylation state of tRNA for feedback on expression of amino acid biosynthetic operons and measuring the flux balance between amino acid synthesis and consumption in protein synthesis (Elf, 2004). Hda can possibly convert DnaA into its ADP-form with a rate proportional to the chromosome copy number since its activity is dependent on active replication forks (Katayama et al., 1998), the number of which, at least at fast growth, is proportional to the number of chromosomes. We find that the *hda* deletion mutant has clear problems controlling initiation, especially at fast growth, where the cells die.

When it comes to conversion to the ATP-form, this could be solved by a simple first-order dissociation of ADP and exchange by ATP. This implies a larger increase in the number of DnaA-ATP molecules in larger volumes. If initiation is triggered at a fixed DnaA-ATP concentration, this mechanism would, however, become very sensitive to the total concentration of DnaA, which goes against our observations. Alternative solutions would be that initiation is responsive to the DnaA-ATP/ADP ratio, or the existence of a saturable catalyst mediating the ADP-to-ATP conversion, with a total activity proportional to the cell volume. The acidic phospholipids may, for example, have this role. Both mechanisms require additional investigation.

In addition to the need of being saturable with respect to DnaA-ADP, there is another caveat with the hypothetical DnaA-ATP/ADP conversion system to monitor the number of chromosomes to volume; it cannot be too efficient or accurate, or we would see a sizer phenotype and not the observed 0.5 correlations between initiations; *i*.*e*. there is clearly memory in the system, which could, for example, be due to fluctuations in the enzyme systems making the conversion on a time scale of the cell cycle.

Finally, we note that RIDA cannot be the main regulator under slow growth conditions where replication is triggered a long time after termination. Most likely, another regulatory scheme is operating in this regime (Berger and ten Wolde, 2021). This mechanism could depend on DnaA-ATP accumulation through gene expression and titration against DnaA boxes or the number of *datA* loci. To speculate more, it may be that a simple initiator-accumulation model based on DnaA expression and binding to DnaA boxes was the primordial control system for slow-growing ancestors of *E. coli* and that the DnaA-ADP/ATP cycling based on RIDA is an adaptation allowing for overlapping replication cycles at fast growth rates.

## Materials and Methods

### Strains and media

The strain *E. coli* MG1655 BW25993 *phi80*-*rph+* (EL544) and derivatives thereof were used in all experiments. The genotypes of all strains are listed in Table S3. If not stated otherwise, we have performed our experiments in M9 minimal medium supplemented with 0.4% succinate and 1× RPMI 1640 amino acid solution (Sigma). All other media used are found in Table S1.

#### Chromosomal modifications

For the construction of chromosomal mutations (deletions, fusions, replacements), λ Red recombineering (Datsenko and Wanner, 2000; Yu et al., 2000) was used. The λ Red genes were expressed from plasmid pSIM5-Tet (Koskiniemi et al., 2011). As selectable/counter selectable markers, we used *Acatsac1* (GenBank acc. # MF124798.1) or *Akansac1*, carrying a *cat* gene (conferring chloramphenicol resistance) or *kan* gene (conferring kanamycin resistance) for positive selection, and the *B. subtilis sacB* gene (conferring sensitivity to sucrose) for negative selection. Generalized transduction using phage P1 *vir* (Thomason et al., 2007) was used for transferring mutations between strains. For many experiments, the chromosomal construct *cobA*::*seqA-venus* was introduced into the strains, keeping the native *seqA* gene intact. In contrast to having *seqA-venus* as the only source of SeqA in the strains, this mutant had no effect on exponential growth.

The chromosomal knockouts of replication regulators (*DARS1, DARS2*, and *datA*) are all marker free and scar free and were made using two different methods: (i) through single strand λ Red, where we cured an inserted selectable/counterselectable marker by an oligo covering the junction of the desired deletion. (ii) by employing the DIRex method (Näsvall, 2017). Mutations were transferred between strains through generalized transduction with phage P1. *hda* was deleted through λ Red recombineering of a *kanR* cassette that exchanged the *hda* reading frame. Recovery and selection were done in M9 succinate medium. All of the other mutant strains were constructed in LB medium.

*Constitutively expressed DnaA mutants*. Firstly, we exchanged an *Acatsac1* cassette in the selectively neutral *galK* locus by PCR-fragments containing *P*_*J23100*_-*dnaA, P*_*J23101b*_-*dnaA*, and *P*_*J23106*_-*dnaA*, selecting sucrose resistant recombinants. Next, the native *dnaA* gene was deleted by replacing it with *Acatsac1*, thereafter *Acatsac1* in turn was replaced by an oligo containing the J23106 promoter (to allow constitutive expression of the rest of the operon). 123 nt were kept at the end of *dnaA*.

*IPTG inducible DnaA*. Two overlapping DNA fragments were simultaneously transformed through electroporation into a strain that expressed the λ Red genes. One fragment was synthesized and contained bp 150–390 of the *lacZ* gene followed by stop codons in all frames, and the *dnaA* gene (including its ribosome binding site) was transcriptionally fused to *mCerulean* MP. The other was a PCR-fragment containing 40 bp homology to the first fragment, a *kan* gene flanked on both sides by Flp recombinase target (FRT) sites, and 40 bp homology to the sequence downstream of the *lacA* gene. The resulting recombinants thus had everything between bp 390 in *lacZ* until just after the *lacA* gene replaced by *dnaA-mCerulean-FRT-kan-FRT*, leaving *dnaA* transcriptionally fused to the beginning of the *lacZ* gene. This construct was transduced to our *seqA-venus* strain and, thereafter, *kan* was removed by expression of Flp recombinase from plasmid pCP20 (Cherepanov and Wackernagel, 1995), leaving a scar containing a single FRT site. Finally, the native *dnaA* gene was replaced by the J23106 promoter as described above. The resultant strain is hence dependent on IPTG induction of *P*_*lac*_*-dnaA* for growth. Similarly to the constitutively expressed DnaA mutants, 123 nt were kept at the end of *dnaA*.

CRISPRi/*dCas9*. The system was adapted from Camsund et al., (2020). *pguide7* plasmids were constructed through Gibson Assembly (Gibson et al., 2009), with the following guide RNA’s: *pgsA* (GGAACAGTGTAAGCAACGTA) and *lacO* (control) (TCCGCTCACAATTCCACATG).

*parS/parB terminus tag*. This system was adapted from Nielsen et al., (2006) and was recently modified by Wiktor et al., (2021). Here, we transformed *parS* into a neutral position in the terminus region by exchanging y*dbL* for *parS*-FRT-*cat*-FRT. *mCherry*-*ParBMt1* (Δ*gtrA*::*p58-mCherry-parB-SpR*; Wiktor et al., 2021) was introduced through P1 transduction.

Other chromosomal constructs originated from other labs and were P1 transduced into our strains (*kanR*-*ypet*-*dnaN* and *dnaQ*-*ypet*-FRT-*kanR*-FRT; Reyes-Lamothe et al., 2010), or introduced in another chromosomal location through λ Red *seqA*-*venus*-FRT-*cat-*FRT (Babic et al., 2008).

### Proteomic analysis with masspectometry

The proteomic analyses were performed as described in Knöppel et al., (2020). Briefly, cultures were started by inoculating a 50 ml succinate medium with 250 *μ*l overnight culture. The cultures were allowed to grow to OD_600_ = 0.2, pelleted by centrifugation, and washed twice in PBS. The pellets were frozen before further preparation. In the medium swap experiment, two flasks with 10 ml succinate medium with 1 mM IPTG were inoculated with 250 *μ*l overnight culture, and the cultures were grown for 3 h, after which they were pelleted by centrifugation and washed twice with succinate medium (without IPTG). The pellets were resuspended in 50 ml pre-warmed succinate medium and allowed to continue growing for 90 or 180 minutes before being pelleted, washed twice in PBS, and frozen. The proteomics core facility at the Sahlgrenska Academy, Gothenburg University, homogenized the samples, digested with trypsin and labeled the peptides with TMT 10-plex isobaric tagging reagents, and performed relative quantification of peptides by LC-MS/MS, as described in Knöppel et al., (2020).

### Microscopy

#### Optical configurations

Five different optical configurations were used, as described below. Table S4 lists which experiments were performed with which microscope configuration.

Configuration 1: An inverted TiE (Nikon) microscope with a CFI Plan Apo lambda 1.45/100x oil objective (Nikon) was used. Phase-contrast images were acquired on a DMK 38UX30 (The Imaging Source) camera. Bright-field and fluorescence images were acquired on an iXon Ultra 888 (Andor) camera. TLED+ (Sutter Instruments) with a white LED was used as a light source for phase-contrast and bright-field images. For fluorescence images, the cells were episcopically illuminated by a 5.3 W/cm^2^ light beam from a 514 nm Genesis CX-STM 2000 mW (Coherent) laser and, when applicable, from a 445 nm 06-MLD 120 mW (Cobolt) laser. The 514 nm laser light was reflected onto the sample using the dichroic mirror Di02-R514 (Semrock) and the emitted fluorescence passed through the same dichroic and was filtered through a BrightLine FF01-474/27 (Semrock) emission filter. The 445 nm laser light was reflected onto the sample using the dichroic mirror Di02-R442 (Semrock) and the emitted fluorescence passed through the same dichroic and was filtered through a BrightLine FF01-550/49 (Semrock) emission filter.

Configuration 2: Similar to Configuration 1, but a red 630 nm LED (Sutter Instruments) was used for phase-contrast, a 600 nm short-pass filter (Edmund Optics) was placed in the microscope camera port for the iXon Ultra, the laser lights were mirrored on FF468/526/596-Di01 (Semrock), and the emitted fluorescence was filtered through a FF01-484/543/702 BrightLine (Semrock) emission filter.

Configuration 3: Similar to Configuration 1, except a 515 nm 06-MLD (Cobolt) laser was used.

Configuration 4: Similar to Configuration 1 but the lasers used for fluorescence imaging were 515 nm (Fandango 150, Cobolt) and 580 nm (VFL, MBP Communications). The laser power was set to 5 W/cm^2^ for both lasers. Fluorescence images were acquired using a Kinetix sCMOS (Teledyne Photometrics) camera. The laser lights were reflected onto the sample using a FF444/521/608-Di01 (Semrock) triple-band dichroic mirror, and the emitted fluorescence passed through the same dichroic mirror. The emitted fluorescence was also passed through a BrightLine FF580-FDi02-T3 (Semrock) dichroic beamsplitter. The split fluorescence was then filtered through BrightLine FF01-505/119-25 (Semrock) and BrightLine FF02-641/75-25 (Semrock) filters and focused on two different parts of the sCMOS camera chip. Phase-contrast images were acquired using a DMK 38UX304 (The Imaging Source) camera. A TLED+ (Sutter Instruments) with a 480 nm LED was used as a light source for the acquisition of phase-contrast images. The transmitted light was passed through a FF444/521/608-Di01 (Semrock) triple-band dichroic mirror and reflected onto the camera using a Di02-R514 (Semrock) dichroic mirror.

Configuration 5: Same as in Wiktor et al., (2021). Three different settings for the Spectra Gen 3 (Lumencor) were used. TEAL was used for YPet-DnaN with 6% power, YELLOW was used for ParB-mCherry with 4.5% power, and BLUE was used for *dnaA-mCerulean* with 6.7% power, unless otherwise stated.

#### Cell preparation

The day before each experiment, strains were inoculated into tubes containing the intended media for the experiment from frozen stock cultures stored at -80 °C. Cells were grown overnight in a 30 or 37 °C shaking incubator (200 rpm) depending on which temperature the experiment was performed at. On the day of the experiment, cells were diluted either 1:100, 1:250, or 1:1000 in fresh medium and grown for 2–4 h before being loaded onto the microfluidic chip, or cells were loaded from stationary phase into the microfluidic chip. Cells were allowed to grow for 6–9 h before starting the experiment if loaded directly from stationary phase and for 2–9 h if the cultures were diluted. For the experiment performed in M9 acetate, cells were first inoculated from -80 °C into LB and grown overnight at 37 °C in a 200 rpm shaking incubator. The next day, cells were diluted 1:100 into fresh M9 acetate medium and grown for roughly 24 h before being loaded onto the chip. The cells were then allowed to grow for roughly 24 hours before image acquisition started. The medium was supplemented with the starting concentrations of IPTG for the experiments using the *P*_*lac*_*-dnaA-mCerulean* and *P*_*lac*_*-dnaA-mCerulean ΔDARS1 ΔDARS2 ΔdatA* strains.

#### Microfluidic experiments

Experiments were performed using a PDMS mother-machine type chip with open-ended channels that allowed for the loading of two separate strains (Baltekin et al., 2017). To keep the medium flowing, pressure on the different ports was maintained with an OB1 MK3+ microfluidic flow controller (Elveflow). This controller was also used to load cells. The microfluidic chips used had four different trap sizes (875, 1000, 1125, and 1250 nm). Table S4 contains which experiments used what trap size. Unless noted, phase-contrast images were acquired every 30 s (100 ms exposure time) and fluorescence images every 2 min (300 ms exposure time).

The duration of the experiments with different carbon sources (Figure 1) was either 9 h (RDM), 24 h (acetate) or 12 h (all other carbon sources). For the RDM and acetate experiments, phase-contrast images were acquired every 20 s and every 2 min, respectively. Fluorescence images were acquired every 80 s and every 8 min for the RDM and acetate experiments, respectively.

The experiments with constitutively expressed DnaA (Figures 2A and S2) and DnaA-ATP/ADP regulatory mutants (Figures 5A and S5A–C) were run for 12–16 h. For the experiments run on Configurations 1 and 3, bright-field images were acquired every 2 min with an exposure time of 100 ms. For the experiments performed on Configuration 2, a bright-field and a corresponding phase-contrast image were acquired before experiments were started. Bright-field images were used for landmark-based registration.

The experiment with different expression levels when *dnaA* was under the control of the *lac* promoter was divided into parts depending on the IPTG concentration (50, 75, 100, 125, or 150 *µ*M; Figures 2B and 2C). Image acquisition was performed for 6 h at each IPTG concentration except for the 150 to 50 *µ*M part, which ran for 10 h. A roughly 15-minute break was taken before swapping from 125 to 150 *µ*M IPTG. To swap between the five different media, an M-switch and a 2-Switch from Fluigent were used. Growth and replication were monitored by imaging a set of cells in phase-contrast every 1 min while bright-field (100 ms exposure time) and 514 nm fluorescence images were acquired every 2 min. The change in mCerulean fluorescence was monitored by imaging a different set of cells. Phase-contrast, bright-field (100 ms exposure time) and 445 nm fluorescence images (300 ms exposure time) were acquired every 16 min. The power of the 445 nm laser was set at 2.97 W/cm^2^.

The duration of the experiments where DnaA expression was turned off was 5.5–6 h (Figures 3, 5B–G and S3), but divided into two parts. The first part lasted for 90 min, and the M9 succinate medium was supplemented with 1 mM IPTG. To change to medium without IPTG, new tubing and connectors were used. This required detaching the used tubing and connectors, which was done while medium was still flowing over the cells. Image acquisition was restarted 16–23 min after the first round of imaging finished, and images were then acquired for 4 h. To follow replication, phase-contrast (80 ms exposure time) and fluorescence images (300 ms exposure time) were acquired every 1 and 2 min, respectively. To measure DnaA expression, cells were imaged every 10 min in phase-contrast (130 ms exposure time) and in fluorescence (100 ms exposure time). Here, a new set of cells were imaged each time to avoid any effect of photobleaching.

The experiment *P*_*lac*_*-dnaA-mCerulean ΔDARS1 ΔDARS2 ΔdatA* with constant DnaA expression (induction with 100 *µ*M IPTG) was run for 12 h (Figure 5H). Imaging was done similarly to the experiments where expression was turned off, except images to measure DnaA expression were acquired every 30 min and the power of the BLUE setting on the light source was set to 18%.

For the *pgsA* repression experiments (Figures 6 and S6), the medium was supplemented with 50 *µ*g/ml kanamycin at all times. Cells were loaded into a chip with succinate medium that had been further supplemented with 0, 0.5 or 1 ng/ml anhydrotetracycline (aTc).

#### Experiments were run for 8 h

The *Δhda* experiments (Figures 4A, 4B, S4E and S4F) were first performed in succinate for 8 h. The medium was then swapped manually to either 0.5× LB or RDM. The swapping procedure was the same as for the experiments where DnaA was turned off. Imaging was performed for 8 h with 0.5× LB and 12 h for RDM.

In the replisome-*ter* distance experiments (Figure S4D), imaging was performed for 10 h. Phase-contrast images were acquired every 1 min (50 ms exposure time). Fluorescence images were acquired every 1 min (150 ms exposure time), with each acquisition triggering both 580 nm and 515 nm lasers back-to-back by the camera through function generators (Tektronix), one for each laser.

### Image analysis

#### Image analysis pipeline

A fully automated image analysis pipeline previously described (Camsund et al., 2020; Wiktor et al., 2021) primarily written in MATLAB (Mathworks) was used. Cell segmentation was performed either with per object ellipse (POE) fit (Ranefall et al., 2016) or nested Unet neural networks (Ronneberger et al., 2015). For the microscopy configurations 1, 2, 3, and 4, landmark based registration was performed between the two cameras. On Configuration 5, landmark based registration was performed in the replisome-*terminus* experiments between the two different emission filters. To do this, 500 nm fluorescent beads (TetraSpeck, Thermo Fisher) were imaged in both channels. Cell-tracking was performed using the Baxter algorithm (Magnusson et al., 2015). The wavelet algorithm was used to detect fluorescent foci (Olivo-Marin, 2002).

#### Post-processing

Custom written MATLAB 2020b (Mathworks) functions and scripts were used for processing the pipeline output. For estimation of single-cell initiation and termination areas, detected foci of either SeqA-Venus or YPet-DnaN were linked together and tracked using the u-track algorithm (Jaqaman et al., 2008). The parameters were optimized to be able to identify the start of tracks by allowing them to merge and split. To avoid truncation issues when tracking dots, cell lineages consisting of either three, four, or five generations were created. Furthermore, these generations were concatenated into a “super-cell” where the area *A*(*t*) at a given time point *t* is given by *A*(*t*) = *A*(*last*) + 2^*i*^(*a*^*c*^(*t*) − *d*_*frac*_*a*^*p*^(*last*)), where *A*(*last*) is the area of the super-cell before division, *i* is an index for the generation in the lineage, *a*^*c*^ are the areas of the cell in generation *i, d*_*frac*_, is the area fraction of the two sisters right after division and *a*^*p*^(*last*) is the area right before division in generation *i* − 1. To create a super-cell, each cell, the cell’s parent, daughters and sister have to be tracked for a certain number of phase-contrast frames (Table S4). Initiation events were defined as the start of a track at least 9 (acetate) or 11 (all other conditions) frames long. Additionally, initiations were not allowed to start on the first two frames in a super-cell and the two adjacent frames to a possible event had to have good segmentation (no gap in cell-tracking, a sufficiently high Jaccard index, and no convergence between cells). A further set of criteria were used to define termination events. Here, tracks could not end on the last two frames of a super-cell. Additionally, tracks had to be located within the boundary of the daughter or in cases where initiation occurred close to division, the long axis coordinate of the last frame of a trajectory could maximally move 5 pixels from mid-cell of the current generation. Only super-cells that had at least one initiation and termination event and contained unique individual cells except the parent generation were used.

Average initiation areas and coefficients of variations (CV) were determined by using MATLAB’s fit(‘gauss’) function where the number of Gaussians corresponded to the number of generations used to generate super-cells. To avoid truncation errors, peaks far away from the edges of super-cells were chosen. Initiation areas in the multi-generational fork plots were calculated as the median of each generation’s initiation distribution. In the *pgsA* repression experiments (Figures 6 and S6A), the average initiation area in bulk was determined the same way as in Camsund et al., (2020).

To get initiation-initiation, initiation-termination, and termination-initiation correlations, all of the events in each super-cell were matched with one another, resulting in data points that could be clustered using a Gaussian mixture model. Then, clusters corresponding to each type of correlation were chosen (Figure S4B). Each cluster was fitted to bivariate Gaussian functions. To avoid truncation bias, a cluster that did not correspond to the edges of the super-cells was chosen. Furthermore, cells in the super-cell structures were all classified as being well segmented (no gaps in cell-tracking, a high enough Jaccard index, and that cells did not converge).

Multi-generational fork plots were created based on pooled super-cells where the cell long axis position of SeqA-Venus or YPet-DnaN is displayed. The long axis offsets were calculated as the average offset from each super-cell. Additionally, in the experiments where *dnaA* expression was turned off, only the mother cells were included in the analysis (cells stuck to the constriction in each channel). Each mother cell had to be tracked for at least 5 minutes to be included in a super-cell structure.

In the *P*_*lac*_*-dnaA-mCerulean* experiments, the mCerulean fluorescence was quantified by first averaging the pixel intensity within the segmentation mask of each cell. Then, a total average for each position was calculated. Background subtraction was performed using a reference strain lacking mCerulean and this was done using positions on the microfluidic chip opposite of each other. Fluorescence half-life was determined by fitting a single exponential to data points ranging from 20 minutes after the media swap until the mCerulean signal plateaued. To calculate the fluorescence fold change, the data points within the fork plot bins were averaged before being normalized, where the highest average signal was defined as 1.

Binned fork plots were created by sorting fluorescent foci based on their detection time relative to the start of the experiment. To calculate average generation times for each bin, the area expansion of all cells within each bin was used. However, if a cell was tracked for fewer than three frames in a bin, the data points were moved into the bin closest in time. Birth and division areas were sorted in the bins where cells were born and divided.

To estimate replisome-*ter* distances, detected foci of YPet-DnaN and ParB-mCherry were tracked separately using the u-track algorithm (Jaqaman et al., 2008), as described above. Imaging of 100 nm fluorescent beads (TetraSpeck, Thermo Fisher) was performed in both of the fluorescence channels imaged on two different parts of the camera chip for landmark-based registration. Based on this registration, fluorescent foci from the same cell detected in the two channels could be paired. Distances were estimated only between paired YPet-DnaN and ParB-mCherry foci that were the closest to each other. Additionally, each YPet-DnaN focus was only paired with one other ParB-mCherry focus for distance estimation. The tracking of YPet-DnaN was used to determine initiation and termination events. The termination events detected for each cell were used as a reference point to compare distances estimated over time in different cells (Figure S4D).

### Determination of *oriC/ter* ratio (NGS approach)

Overnight cultures of EL544 were grown in 1 ml medium (LB, succinate, and glucose) at 37 °C. The cultures were diluted at least 250-fold in 25 ml fresh media pre-heated to 30 or 37 °C and grown to OD_600_ = 0.2 (LB 30 and 37 °C), 0.17 (succinate 37 °C), and 0.14 (succinate and glucose 30 °C) where after rifampicin was added to the final concentration of 0.3 mg/ml and any ongoing transcription was allowed to terminate by leaving the cultures in the shaking incubator for 2 min before withdrawing 2× 1.5 ml samples. The cells in the withdrawn samples were quickly pelleted at room temperature and frozen at -84 °C (*i*.*e*. the cells were frozen within 4 min after adding rifampicin). In addition, 0.5 ml of a sample with stationary cells grown in LB at 37 °C for 1.5 days was pelleted and frozen. DNA was prepared using the MasterPure™ Complete DNA and RNA purification kit (Lucigen), according to the manufacturer’s instructions. Illumina libraries were prepared using the TruSeq DNA PCR-Free library kit and the samples were sequenced using one lane of a MiSeq run. TrimGalore (Krueger et al., 2021) was used to remove adapter sequences and Bowtie2 (Langmead and Salzberg, 2012) to map the sequencing reads over the reference genome of EL544. The genome position of the first base of each mapped read was binned into either ∼30 kb or ∼1.5 kb large bins (Figure S4A).

### Quantitative PCR

Experiment presented in Figure S3A: overnight cultures were grown in 1 ml medium supplemented with 1 mM IPTG (duplicates for wt and triplicates for the *P*_*lac*_*-dnaA-mCerulean strain*). The cultures were diluted to 1:100 in 25 ml fresh pre-heated media supplemented with 1 mM IPTG and grown to OD_600_ = 0.2 after which samples for downstream RNA extraction were withdrawn (t_0_). To mimic the medium swap in the fluidics experiment, the cells in the remainder of the cultures were carefully washed four times through centrifugation in media without IPTG. Then the cells were resuspended in 100 ml pre-heated medium and grown for four hours at 30 °C with shaking. Samples were withdrawn after 20, 40, 60, 120, and 240 min (*i*.*e*. t_1_, t_2_, t_3_, t_4_ and t_5_).

Experiment presented in Figure S3B: quadruplicate overnight cultures were grown in 1 ml medium at varying IPTG concentrations (*P*_*lac*_*-dnaA-mCerulean strain*: 1, 0.05, and 0.005 mM). The cultures were diluted to 1:1000 in 50 ml fresh pre-heated media supplemented with the same IPTG concentrations and grown to OD_600_ ≈ 0.2 when samples for downstream RNA extraction were withdrawn. However, the sample with 0.005 mM IPTG never seemed to reach OD_600_ = 0.2, although the overnight culture at the same concentration was fully grown. It was instead harvested after about 5 h. The samples with 0 mM IPTG were taken from the 1 mM cultures and treated similarly to the experiment presented in Figure S3A, although only one time point after the swap was taken (240 min).

RNA from both experiments was prepared using the PureLink RNA Mini Kit (ThermoFisher) and DNase treated using the Turbo DNA-free kit (Invitrogen). The RNA was reverse-transcribed into cDNA through the High Capacity Reverse Transcription kit (ThermoFisher), and the Power SYBR Green PCR mix (ThermoFisher) was used for the quantitative PCR reactions according to the manufacturer’s instructions. Quantitative PCRs were performed in technical replicates on all samples, using primer pairs for *dnaA* and the two reference genes, *cysG* and *hsaT*. After removing data from strange looking flat curves, the technical replicates were averaged and the fold change was calculated according to the 2^−ΔΔ*CT*^ method (Kenneth et al., 2001). The geometrical mean of the two reference genes was used in the calculations. Primers used in RT-qPCR were *hcaT_f/r* and *cycG_f/r* (Zhou et al., 2011) *ori_3_f/r* and *ter_14_f/r* (Fernández-Coll et al., 2020), and *dnaA_f/r* (Table S5). The samples used in the qPCR experiment presented in Figure S6A were the same as the ones prepared for the NGS Marker Frequency analysis in the same figure. Quantitative PCRs were performed with 0.1 and 1 *µ*l template and in technical duplicates for primer pair *ori 3* and single PCRs for primer pair *ter 14*. The technical replicates were averaged and the *ori*/*ter* ratio was calculated according to the 2^−ΔΔ*CT*^ method (Kenneth et al., 2001). Primers used in qPCR are found in Table S5.

### Rifampicin (rif) run-out experiments

Overnight cultures (made in quadruplicates for the experiment in Figure S2A and duplicates for Figure S5D) were diluted to grow for at least 10 generations in a shaking incubator before reaching OD_600_ ≈ 0.05. To stop initiation and division, rifampicin and cephalexin were added to the final concentrations of 0.3 mg/ml and 0.01 mg/ml, respectively (from rifampicin stock 30 mg/ml in DMSO and cephalexin stock 10 mg/ml in H_2_O). Since any ongoing replication is still active after adding the antibiotics, the replication forks will go on until termination, allowing chromosome copy number to serve as a measurement of *oriC* number at the time point of adding the antibiotics. After 4–5 h of further incubation in the shaking incubator, 5 ml of the samples were fixed in 50 ml ice cold 70% EtOH and stored at -20 °C.

For flow cytometry analysis, cells in 1 ml aliquots of the stored samples were spun down and re-suspended in 1 ml of 50 mM Tris-Mg (pH 7.5) buffer supplemented with 10 mM MgSO_4_. The cells were concentrated through centrifugation, and 0.8 ml of the supernatant was removed. 1–2 *µ*l SYTOX green was added to stain the DNA in the cells, and the DNA content in the fixed cells was analyzed with a MACSquant Analyzer Flow Cytometer.

### Growth rate determinations in plate reader

The measurements were performed by diluting overnight cultures grown in succinate medium 1:1000 in fresh medium and, thereafter, the increase in optical density (OD_600_) over time was measured using a Bioscreen C Reader (Oy Growth Curves) with shaking. Growth rates during the exponential phase were calculated and normalized to the growth of a reference strain included in the same experiment. For the experiment presented in Figure S3C, the overnight cultures were grown in succinate medium supplemented with 0.01 mM IPTG and diluted in fresh media containing 1, 0.1, 0.01, or 0 mM IPTG.

### Growth rate determinations in E-flasks

#### Deletion of hda

Overnight cultures in duplicates of EL3408 (Δ*hda*) and EL562 (reference strain) were grown in succinate medium at 37 °C. The cultures were diluted 1:5000 while OD_600_ was monitored until ≈ 0.02 with a Visible Spectrophotometer, PV4 (VWR). At this point, the medium was swapped to 0.5× LB by diluting 1:50 into pre-warmed 0.5× LB medium. At OD_600_ ≈ 0.13 the reference cultures were re-diluted 1:50 in 0.5× LB medium. To test that the residual succinate medium did not affect growth after medium swap, we filtered the remaining succinate medium over 0.2 *μ*m Filtropur V25 (Sarstedt) filters and rinsed once with pre-heated 0.5× LB medium, where after the cells were resuspended in 0.5× LB medium to the starting volume and inoculated at ≈ 1:50 in 0.5× LB medium. The growth of the filtered cultures was indistinguishable from the non-filtered (data not shown). During analysis, we corrected for dilution factors.

#### Repression of pgsA

Overnight cultures of EL3242 (*P*_*J23106*_*-dnaA lacO* gRNA), EL3244 (*P*_*J23106*_*-dnaA pgsA* gRNA) EL3298 (*P*_*J23106*_*-dnaA lacO* gRNA Δ*DARS1* Δ*DARS2*), EL3299, were (*P*_*J23106*_*-dnaA pgsA* gRNA Δ*DARS1* Δ*DARS2*) were grown in succinate medium supplemented with 50 *µ*g/ml Kan at 30 °C. Cultures were diluted at least 1:20000 into pre-warmed medium ±1 ng/ml aTc. The cultures were allowed to grow for ∼16 h before re-dilution into the same media (pre-warmed) to a final OD_600_ of about 0.005 and OD_600_ was monitored over time. The experiment was repeated three times.

## Acknowledgements

We would like to thank our colleagues Jimmy Larsson, who performed some of the microfluidic experiments, and Spartak Zikrin, who worked on the analysis pipeline. We also thank Irmeli Barkefors and Joakim Näsvall for critical reading of the manuscript, and Pieter Rein ten Wolde and Suckjoon Jun for constructive discussions. Rodrigo Reyes Lamothe provided us with strains containing the YPet-DnaN and the DnaQ-YPet constructs. The Proteomics Core Facility at Sahlgrenska Academy, Gothenburg University, performed the analysis for protein quantification, and SciLifeLab in Uppsala performed the Marker Frequency Illumina sequencing. This study was made possible by grants from the ERC (Advanced grant no. 885360), the Swedish Research Council (grant no. 2016-06213 and 2018-03958), the Knut and Alice Wallenberg Foundation (grant no. 2016.0077, 2017.0291 and 2019.0439), and the eSSENCE e-science initiative. The computations and data management were enabled by resources provided by the Swedish National Infrastructure for Computing (SNIC) at UPPMAX, partially funded by the Swedish Research Council through grant agreement no. 2018-05973.

