## Supplemental Table S4 for "The coordination of replication initiation with growth rate in *Escherichia coli*"

| Figure | # cells | # births | # divisions | # initiations | # lineages | # dot distances | Confiuration # | Chip size | Segmentatio | Minimum cell track length (mir | Minimum parent cell track length ( | Minimum daughter cell track length (n | Minimum sister cell track length (m | Number of repeats |
|---|---|---|---|---|---|---|---|---|---|---|---|---|---|---|
| 1B | NA | NA | NA | NA | 1484 | NA | 5 | 1000 | Unet | 10 | 5 | 5 | 5 | NA |
| 1C (acetate no AA) | 14095 | NA | NA | NA | NA | NA | 5 | 875 | Unet | 40 | 20 | 20 | NA | 1 |
| 1C (lactose no AA) | 23112 | NA | NA | NA | NA | NA | 5 | 875 | Unet | 10 | 5 | 5 | NA | 2 |
| 1C (lactose low AA) | 34728 | NA | NA | NA | NA | NA | 5 | 1000 | Unet | 10 | 5 | 5 | NA | 3 |
| 1C (succinate) | 39297 | NA | NA | NA | NA | NA | 5 | 1000 | Unet | 10 | 5 | 5 | NA | 2 |
| 1C (mannose) | 21092 | NA | NA | NA | NA | NA | 5 | 1000 | Unet | 10 | 5 | 5 | NA | 2 |
| 1C (gluconate) | 28670 | NA | NA | NA | NA | NA | 5 | 1125 | Unet | 10 | 5 | 5 | NA | 1 |
| 1C (glucose) | 14785 | NA | NA | NA | NA | NA | 5 | 1250 | Unet | 10 | 5 | 5 | NA | 3 |
| 1C (RDM) | 11078 | NA | NA | NA | NA | NA | 5 | 1250 | Unet | 6.67 | 3.33 | 3.33 | NA | 1 |
| 1D | 88624[a] | NA | NA | NA | NA | NA | 5 | NA | Unet | NA | NA | NA | NA | NA |
| 2A ($P_{wt}$-*dnaA*) | 8348 | NA | NA | NA | NA | NA | 3 | 1000 | POE | 10 | 5 | 5 | NA | 26 |
| 2A ($P_{J23106}$-*dnaA*) | 7983 | NA | NA | NA | NA | NA | 3 | 1000 | POE | 10 | 5 | 5 | NA | 19 |
| 2A ($P_{J23100}$-*dnaA*) | 6391 | NA | NA | NA | NA | NA | 2 | 1000 | POE | 10 | 5 | 5 | NA | 3 |
| 2A ($P_{J23101b}$-*dnaA*) | 10412 | NA | NA | NA | NA | NA | 1 | 1000 | POE | 10 | 5 | 5 | NA | 3 |
| 2B (50 $\mu$ M) | NA | 812 | 889 | NA | NA | NA | 1 | 1000 | POE | 10 | 5 | 5 | 5 | 1 |
| 2B (75 $\mu$ M) | NA | 570 | 607 | NA | NA | NA | 1 | 1000 | POE | 10 | 5 | 5 | 5 | 1 |
| 2B (100 $\mu$ M) | NA | 688 | 674 | NA | NA | NA | 1 | 1000 | POE | 10 | 5 | 5 | 5 | 1 |
| 2B (125 $\mu$ M) | NA | 639 | 890 | NA | NA | NA | 1 | 1000 | POE | 10 | 5 | 5 | 5 | 1 |
| 2B (150 $\mu$ M) | NA | 839 | 890 | NA | NA | NA | 1 | 1000 | POE | 10 | 5 | 5 | 5 | 1 |
| 2B (50 $\mu$ M) | NA | 868 | 864 | NA | NA | NA | 1 | 1000 | POE | 10 | 5 | 5 | 5 | 1 |
| 3A (0–90 min) | NA | 1609 | 1609 | NA | NA | NA | 5 | 1000 | Unet | 5 | 3 | 3 | 3 | 3 |
| 3A (111–141 min) | NA | 1128 | 4 | NA | NA | NA | 5 | 1125 | Unet | 5 | 3 | 3 | 3 | 3 |
| 3A (141–171 min) | NA | 1364 | 241 | NA | NA | NA | 5 | 1125 | Unet | 5 | 3 | 3 | 3 | 3 |
| 3A (171–201 min) | NA | 1562 | 1377 | NA | NA | NA | 5 | 1125 | Unet | 5 | 3 | 3 | 3 | 3 |
| 3A (201–231 min) | NA | 1360 | 1491 | NA | NA | NA | 5 | 1125 | Unet | 5 | 3 | 3 | 3 | 3 |
| 3A (231–261 min) | NA | 1355 | 1384 | NA | NA | NA | 5 | 1125 | Unet | 5 | 3 | 3 | 3 | 3 |
| 3A (261–291 min) | NA | 1334 | 1397 | NA | NA | NA | 5 | 1125 | Unet | 5 | 3 | 3 | 3 | 3 |
| 3A (291–321 min) | NA | 478 | 1361 | NA | NA | NA | 5 | 1125 | Unet | 5 | 3 | 3 | 3 | 3 |
| 3A (321–351 min) | NA | 5 | 1331 | NA | NA | NA | 5 | 1125 | Unet | 5 | 3 | 3 | 3 | 3 |
| 3B (0–90 min) | NA | 1063 | 1063 | NA | NA | NA | 5 | 1125 | Unet | 5 | 3 | 3 | 3 | 3 |
| 3B (141–171 min) | NA | 1091 | 299 | NA | NA | NA | 5 | 1125 | Unet | 5 | 3 | 3 | 3 | 3 |
| 3C ($P_{J23106}$-*dnaA*) | NA | NA | NA | NA | 103 | NA | 5 | 1125 | Unet | 5 | 3 | 3 | 3 | 2 |
| 3C ($P_{lac}$-*dnaA-mcerulean*) | NA | NA | NA | NA | 139 | NA | 5 | 1125 | Unet | 5 | 3 | 3 | 3 | 2 |
| 3D ($P_{J23106}$-*dnaA*) | NA | NA | NA | 564 | 103 | NA | 5 | 1125 | Unet | 5 | 3 | 3 | 3 | 2 |
| 3D ($P_{lac}$-*dnaA-mcerulean*) | NA | NA | NA | 559 | 139 | NA | 5 | 1125 | Unet | 5 | 3 | 3 | 3 | 2 |
| 4C (M9 succ wt) | NA | 16031 | 16031 | NA | NA | NA | 2 | 1125 | Unet | 10 | 5 | 5 | NA | 5 |
| 4C (M9 succ Δ*hda*) | NA | 14391 | 14391 | NA | NA | NA | 2 | 1125 | Unet | 10 | 5 | 5 | NA | 5 |
| 4C (0.5× LB wt) | NA | 3479 | 3479 | NA | NA | NA | 2 | 1125 | Unet | 2.5 | 1.5 | 1.5 | NA | 3 |
| 4C (0.5× LB Δ*hda*) | NA | 352 | 550 | NA | NA | NA | 2 | 1125 | Unet | 2.5 | 1.5 | 1.5 | NA | 3 |
| 4C (RDM wt) | NA | 3146 | 3093 | NA | NA | NA | 2 | 1125 | Unet | 2.5 | 1.5 | 1.5 | NA | 2 |
| 4C (RDM Δ*hda*) | NA | 900 | 1203 | NA | NA | NA | 2 | 1125 | Unet | 2.5 | 1.5 | 1.5 | NA | 2 |
| 5A ($P_{wt}$-*dnaA*) | 8348 | NA | NA | NA | NA | NA | 3 | 1000 | POE | 10 | 5 | 5 | NA | 26 |
| 5A ($P_{J23106}$-*dnaA*) | 7983 | NA | NA | NA | NA | NA | 3 | 1000 | POE | 10 | 5 | 5 | NA | FILL IN |
| 5A ($P_{J23106}$-*dnaA* Δ*DARS1*) | 4209 | NA | NA | NA | NA | NA | 3 | 1000 | POE | 10 | 5 | 5 | NA | 2 |
| 5A ($P_{J23106}$-*dnaA* Δ*DARS2*) | 6168 | NA | NA | NA | NA | NA | 1 | 1000 | POE | 10 | 5 | 5 | NA | 2 |
| 5A ($P_{J23106}$-*dnaA* Δ*datA*) | 8087 | NA | NA | NA | NA | NA | 1 | 1000 | POE | 10 | 5 | 5 | NA | 2 |
| 5A ($P_{J23106}$-*dnaA* Δ*DARS12*) | 6385 | NA | NA | NA | NA | NA | 3 | 1000 | POE | 10 | 5 | 5 | NA | 2 |
| 5A ($P_{J23106}$-*dnaA* Δ*DARS1* Δ*datA*) | 4623 | NA | NA | NA | NA | NA | 2 | 1000 | POE | 10 | 5 | 5 | NA | 3 |
| 5A ($P_{J23106}$-*dnaA* Δ*DARS2* Δ*datA*) | 8217 | NA | NA | NA | NA | NA | 3 | 1000 | POE | 10 | 5 | 5 | NA | 2 |
| 5A ($P_{J23106}$-*dnaA* Δ*DARS12* Δ*datA*) | 7792 | NA | NA | NA | NA | NA | 1 | 1000 | POE | 10 | 5 | 5 | NA | 1 |
| 5B (0–90 min) | NA | 1647 | 1647 | NA | NA | NA | 5 | 1125 | Unet | 5 | 3 | 3 | 3 | 4 |
| 5B (106–136 min) | NA | 1276 | 7 | NA | NA | NA | 5 | 1125 | Unet | 5 | 3 | 3 | 3 | 4 |
| 5B (136–166 min) | NA | 1497 | 416 | NA | NA | NA | 5 | 1125 | Unet | 5 | 3 | 3 | 3 | 4 |
| 5B (166–196 min) | NA | 1514 | 1408 | NA | NA | NA | 5 | 1125 | Unet | 5 | 3 | 3 | 3 | 4 |
| 5B (196–226 min) | NA | 1582 | 1491 | NA | NA | NA | 5 | 1125 | Unet | 5 | 3 | 3 | 3 | 4 |
| 5B (226–256 min) | NA | 1231 | 1373 | NA | NA | NA | 5 | 1125 | Unet | 5 | 3 | 3 | 3 | 4 |
| 5B (256–286 min) | NA | 459 | 1409 | NA | NA | NA | 5 | 1125 | Unet | 5 | 3 | 3 | 3 | 4 |
| 5B (286–316 min) | NA | 22 | 859 | NA | NA | NA | 5 | 1125 | Unet | 5 | 3 | 3 | 3 | 4 |
| 5C (0–90 min) | NA | 1349 | 1439 | NA | NA | NA | 5 | 1125 | Unet | 5 | 3 | 3 | 3 | 4 |
| 5C (166–196 min) | NA | 917 | 929 | NA | NA | NA | 5 | 1125 | Unet | 5 | 3 | 3 | 3 | 4 |
| 5D ($P_{J23106}$-*dnaA*) | NA | NA | NA | NA | 70 | NA | 5 | 1125 | Unet | 5 | 3 | 3 | 3 | 4 |
| 5D ($P_{lac}$-*dnaA-mcerulean* Δ*DARS12* Δ*datA*) | NA | NA | NA | NA | 254 | NA | 5 | 1125 | Unet | 5 | 3 | 3 | 3 | 4 |
| 5E ($P_{J23106}$-*dnaA*) | NA | NA | NA | 367 | 70 | NA | 5 | 1125 | Unet | 5 | 3 | 3 | 3 | 4 |
| 5E ($P_{lac}$-*dnaA-mcerulean* Δ*DARS12* Δ*datA*) | NA | NA | NA | 874 | 254 | NA | 5 | 1125 | Unet | 5 | 3 | 3 | 3 | 4 |
| 5H ($P_{J23106}$-*dnaA*) | 12415 | NA | NA | NA | NA | NA | 5 | 1000 | Unet | 10 | 5 | 5 | 5 | 1 |
| 5H ($P_{lac}$-*dnaA-mcerulean* Δ*DARS12* Δ*datA*) | 14723 | NA | NA | NA | NA | NA | 5 | 1000 | Unet | 10 | 5 | 5 | 5 | 1 |
| 6 (0 ng in wt) | 13773 | NA | NA | NA | NA | NA | 2 | 1000 | Unet | 10 | 5 | 5 | NA | 1 |
| 6 (0 ng in Δ*DARS12*) | 7760 | NA | NA | NA | NA | NA | 2 | 1000 | Unet | 10 | 5 | 5 | NA | 1 |
| 6 (1 ng in wt) | 6696 | NA | NA | NA | NA | NA | 2 | 1000 | Unet | 10 | 5 | 5 | NA | 3 |
| 6 (1 ng in Δ*DARS12*) | 4294 | NA | NA | NA | NA | NA | 2 | 1000 | Unet | 10 | 5 | 5 | NA | 3 |
| S1A (SeqA) | 36477 | NA | NA | NA | NA | NA | 5 | 1000 | Unet | 10 | 5 | 5 | NA | 2 |
| S1A (DnaQ) | 3731 | NA | NA | NA | NA | NA | 5 | 1000 | Unet | 10 | 5 | 5 | NA | 3 |
| S1A (DnaN) | 39297 | NA | NA | NA | NA | NA | 5 | 1000 | Unet | 10 | 5 | 5 | NA | 2 |
| S1B (30 °C) | 39297 | NA | NA | NA | NA | NA | 5 | 1000 | Unet | 10 | 5 | 5 | NA | 2 |
| S1B (37 °C) | 8289 | NA | NA | NA | NA | NA | 2 | 1000 | POE | 10 | 5 | 5 | NA | 1 |
| S2B ($P_{wt}$-*dnaA*) | 8348 | NA | NA | NA | NA | NA | 3 | 1000 | POE | 10 | 5 | 5 | NA | 26 |
| S2B ($P_{wt}$-*dnaA*) | 6926 | NA | NA | NA | NA | NA | 2 | 1000 | POE | 10 | 5 | 5 | NA | 26 |
| S2B ($P_{wt}$-*dnaA*) | 11702 | NA | NA | NA | NA | NA | 1 | 1000 | POE | 10 | 5 | 5 | NA | 26 |
| S2B ($P_{J23106}$-*dnaA*) | 7983 | NA | NA | NA | NA | NA | 3 | 1000 | POE | 10 | 5 | 5 | NA | 19 |
| S2B ($P_{J23100}$-*dnaA*) | 6391 | NA | NA | NA | NA | NA | 2 | 1000 | POE | 10 | 5 | 5 | NA | 3 |
| S2B ($P_{J23101b}$-*dnaA*) | 10412 | NA | NA | NA | NA | NA | 1 | 1000 | POE | 10 | 5 | 5 | NA | 3 |
| S2CD ($P_{wt}$-*dnaA*) | 8348 | NA | NA | 3973 | NA | NA | 3 | 1000 | POE | 10 | 5 | 5 | 5 | 26 |
| S2CD ($P_{wt}$-*dnaA*) | 6926 | NA | NA | 2721 | NA | NA | 2 | 1000 | POE | 10 | 5 | 5 | 5 | 26 |
| S2CD ($P_{wt}$-*dnaA*) | 11702 | NA | NA | 7744 | NA | NA | 1 | 1000 | POE | 10 | 5 | 5 | 5 | 26 |
| S2CD ($P_{J23106}$-*dnaA*) | 7983 | NA | NA | 5001 | NA | NA | 3 | 1000 | POE | 10 | 5 | 5 | 5 | 19 |
| S2CD ($P_{J23100}$-*dnaA*) | 6391 | NA | NA | 2999 | NA | NA | 2 | 1000 | POE | 10 | 5 | 5 | 5 | 3 |
| S2CD ($P_{J23101b}$-*dnaA*) | 10412 | NA | NA | 4803 | NA | NA | 1 | 1000 | POE | 10 | 5 | 5 | 5 | 3 |
| S3D (0–90 min) | NA | 1243 | 1196 | NA | NA | NA | 5 | 1000 | Unet | 5 | 3 | 3 | NA | 3 |
| S3D (113–143 min) | NA | 909 | 9 | NA | NA | NA | 5 | 1000 | Unet | 5 | 3 | 3 | NA | 3 |
| S3D (143–173 min) | NA | 884 | 296 | NA | NA | NA | 5 | 1000 | Unet | 5 | 3 | 3 | NA | 3 |
| S3D (173–203 min) | NA | 902 | 927 | NA | NA | NA | 5 | 1000 | Unet | 5 | 3 | 3 | NA | 3 |
| S3D (203–233 min) | NA | 941 | 955 | NA | NA | NA | 5 | 1000 | Unet | 5 | 3 | 3 | NA | 3 |
| S3D (233–263 min) | NA | 823 | 877 | NA | NA | NA | 5 | 1000 | Unet | 5 | 3 | 3 | NA | 3 |
| S3D (263–293 min) | NA | 905 | 875 | NA | NA | NA | 5 | 1000 | Unet | 5 | 3 | 3 | NA | 3 |
| S3D (293–323 min) | NA | 256 | 898 | NA | NA | NA | 5 | 1000 | Unet | 5 | 3 | 3 | NA | 3 |
| S3D (323–353 min) | NA | 1 | 784 | NA | NA | NA | 5 | 1000 | Unet | 5 | 3 | 3 | NA | 3 |
| S3E (0–90 min) | NA | 1049 | 996 | NA | NA | NA | 5 | 1000 | Unet | 5 | 3 | 3 | NA | 3 |
| S3E (143–173 min) | NA | 712 | 241 | NA | NA | NA | 5 | 1000 | Unet | 5 | 3 | 3 | NA | 3 |
| S4C (acetate no AA) | 14095 | NA | NA | NA | NA | NA | 5 | 875 | Unet | 40 | 20 | 20 | NA | 1 |
| S4C (lactose no AA) | 23112 | NA | NA | NA | NA | NA | 5 | 875 | Unet | 10 | 5 | 5 | NA | 2 |
| S4C (lactose low AA) | 34728 | NA | NA | NA | NA | NA | 5 | 1000 | Unet | 10 | 5 | 5 | NA | 3 |
| S4C (succinate) | 39297 | NA | NA | NA | NA | NA | 5 | 1000 | Unet | 10 | 5 | 5 | NA | 2 |
| S4C (mannose) | 21092 | NA | NA | NA | NA | NA | 5 | 1000 | Unet | 10 | 5 | 5 | NA | 2 |
| S4C (gluconate) | 28670 | NA | NA | NA | NA | NA | 5 | 1125 | Unet | 10 | 5 | 5 | NA | 1 |
| S4C (glucose) | 14785 | NA | NA | NA | NA | NA | 5 | 1250 | Unet | 10 | 5 | 5 | NA | 3 |
| S4C (RDM) | 11078 | NA | NA | NA | NA | NA | 5 | 1250 | Unet | 6.67 | 3.33 | 3.33 | NA | 1 |

| | | | | | | | | | | | | | | | |
|---|---|---|---|---|---|---|---|---|---|---|---|---|---|---|---|
| S4D (lactose low AA) | 5955 | NA | NA | NA | NA | 59498 | 4 | 1000 | Unet | 40 | 10 | 3 | 3 | 1 | |
| S4D (mannose) | 8638 | NA | NA | NA | NA | 31524 | 4 | 1000 | Unet | 40 | 10 | 3 | 3 | 1 | |
| S4E ($P_{wt}$-*dnaA;* 0–80 min, LB) | NA | 2168 | 808 | NA | NA | NA | 2 | 1000 | Unet | 2.5 | 1.5 | 1.5 | NA | | 3 |
| S4E ($P_{wt}$-*dnaA;* 80–160 min, LB) | NA | 3481 | 3430 | NA | NA | NA | 2 | 1000 | Unet | 2.5 | 1.5 | 1.5 | NA | | 3 |
| S4E ($P_{wt}$-*dnaA;* 160–240 min, LB) | NA | 3479 | 3479 | NA | NA | NA | 2 | 1000 | Unet | 2.5 | 1.5 | 1.5 | NA | | 3 |
| S4E ($P_{wt}$-*dnaA;* 240–320 min, LB) | NA | 3421 | 3368 | NA | NA | NA | 2 | 1000 | Unet | 2.5 | 1.5 | 1.5 | NA | | 3 |
| S4E ($P_{wt}$-*dnaA;* 320–400 min, LB) | NA | 3575 | 3580 | NA | NA | NA | 2 | 1000 | Unet | 2.5 | 1.5 | 1.5 | NA | | 3 |
| S4E ($P_{wt}$-*dnaA;* 400–480 min, LB) | NA | 1864 | 3323 | NA | NA | NA | 2 | 1000 | Unet | 2.5 | 1.5 | 1.5 | NA | | 3 |
| S4E (*Δhda;* 0–80 min, LB) | NA | 1263 | 520 | NA | NA | NA | 2 | 1000 | Unet | 2.5 | 1.5 | 1.5 | NA | | 3 |
| S4E (*Δhda;* 80–160 min, LB) | NA | 759 | 1100 | NA | NA | NA | 2 | 1000 | Unet | 2.5 | 1.5 | 1.5 | NA | | 3 |
| S4E (*Δhda;* 160–240 min, LB) | NA | 352 | 550 | NA | NA | NA | 2 | 1000 | Unet | 2.5 | 1.5 | 1.5 | NA | | 3 |
| S4E (*Δhda;* 240–320 min, LB) | NA | 177 | 253 | NA | NA | NA | 2 | 1000 | Unet | 2.5 | 1.5 | 1.5 | NA | | 3 |
| S4E (*Δhda;* 320–400 min, LB) | NA | 163 | 190 | NA | NA | NA | 2 | 1000 | Unet | 2.5 | 1.5 | 1.5 | NA | | 3 |
| S4E (*Δhda;* 400–480 min, LB) | NA | 78 | 179 | NA | NA | NA | 2 | 1000 | Unet | 2.5 | 1.5 | 1.5 | NA | | 3 |
| S4F ($P_{wt}$-*dnaA;* 0–80 min, RDM) | NA | 1968 | 666 | NA | NA | NA | 2 | 1000 | Unet | 2.5 | 1.5 | 1.5 | NA | | 2 |
| S4F ($P_{wt}$-*dnaA;* 80–160 min, RDM) | NA | 2627 | 2630 | NA | NA | NA | 2 | 1000 | Unet | 2.5 | 1.5 | 1.5 | NA | | 2 |
| S4F ($P_{wt}$-*dnaA;* 160–240 min, RDM) | NA | 3146 | 3093 | NA | NA | NA | 2 | 1000 | Unet | 2.5 | 1.5 | 1.5 | NA | | 2 |
| S4F ($P_{wt}$-*dnaA;* 240–320 min, RDM) | NA | 3069 | 3069 | NA | NA | NA | 2 | 1000 | Unet | 2.5 | 1.5 | 1.5 | NA | | 2 |
| S4F ($P_{wt}$-*dnaA;* 320–400 min, RDM) | NA | 3067 | 3078 | NA | NA | NA | 2 | 1000 | Unet | 2.5 | 1.5 | 1.5 | NA | | 2 |
| S4F ($P_{wt}$-*dnaA;* 400–480 min, RDM) | NA | 3065 | 3063 | NA | NA | NA | 2 | 1000 | Unet | 2.5 | 1.5 | 1.5 | NA | | 2 |
| S4F ($P_{wt}$-*dnaA;* 480–560 min, RDM) | NA | 3116 | 3093 | NA | NA | NA | 2 | 1000 | Unet | 2.5 | 1.5 | 1.5 | NA | | 2 |
| S4F ($P_{wt}$-*dnaA;* 560–640 min, RDM) | NA | 3060 | 3057 | NA | NA | NA | 2 | 1000 | Unet | 2.5 | 1.5 | 1.5 | NA | | 2 |
| S4F ($P_{wt}$-*dnaA;* 640–720 min, RDM) | NA | 1636 | 3005 | NA | NA | NA | 2 | 1000 | Unet | 2.5 | 1.5 | 1.5 | NA | | 2 |
| S4F (*Δhda;* 0–80 min, RDM) | NA | 1931 | 766 | NA | NA | NA | 2 | 1000 | Unet | 2.5 | 1.5 | 1.5 | NA | | 2 |
| S4F (*Δhda;* 80–160 min, RDM) | NA | 1209 | 1569 | NA | NA | NA | 2 | 1000 | Unet | 2.5 | 1.5 | 1.5 | NA | | 2 |
| S4F (*Δhda;* 160–240 min, RDM) | NA | 900 | 1203 | NA | NA | NA | 2 | 1000 | Unet | 2.5 | 1.5 | 1.5 | NA | | 2 |
| S4F (*Δhda;* 240–320 min, RDM) | NA | 513 | 727 | NA | NA | NA | 2 | 1000 | Unet | 2.5 | 1.5 | 1.5 | NA | | 2 |
| S4F (*Δhda;* 320–400 min, RDM) | NA | 313 | 404 | NA | NA | NA | 2 | 1000 | Unet | 2.5 | 1.5 | 1.5 | NA | | 2 |
| S4F (*Δhda;* 400–480 min, RDM) | NA | 285 | 298 | NA | NA | NA | 2 | 1000 | Unet | 2.5 | 1.5 | 1.5 | NA | | 2 |
| S4F (*Δhda;* 480–560 min, RDM) | NA | 231 | 252 | NA | NA | NA | 2 | 1000 | Unet | 2.5 | 1.5 | 1.5 | NA | | 2 |
| S4F (*Δhda;* 560–640 min, RDM) | NA | 164 | 209 | NA | NA | NA | 2 | 1000 | Unet | 2.5 | 1.5 | 1.5 | NA | | 2 |
| S4F (*Δhda;* 640–720 min, RDM) | NA | 57 | 175 | NA | NA | NA | | 1000 | Unet | 2.5 | 1.5 | 1.5 | NA | | 2 |
| S5A ($P_{wt}$-*dnaA* ; nr 1 from left) | 8348 | NA | NA | NA | NA | NA | 3 | 1000 | POE | 10 | 5 | 5 | NA | 26 | |
| S5A ($P_{wt}$-*dnaA* ; nr 2 from left) | 7759 | NA | NA | NA | NA | NA | 2 | 1000 | POE | 10 | 5 | 5 | NA | 26 | |
| S5A ($P_{wt}$-*dnaA* ; nr 3 from left) | 6066 | NA | NA | NA | NA | NA | 1 | 1000 | POE | 10 | 5 | 5 | NA | 26 | |
| S5A ($P_{wt}$-*dnaA* ; nr 4 from left) | 7212 | NA | NA | NA | NA | NA | 1 | 1000 | POE | 10 | 5 | 5 | NA | 26 | |
| S5A ($P_{wt}$-*dnaA* ; nr 5 from left) | 7924 | NA | NA | NA | NA | NA | 1 | 1000 | POE | 10 | 5 | 5 | NA | 26 | |
| S5A ($P_{J23106}$-*dnaA* ; nr 6 from left) | 7983 | NA | NA | NA | NA | NA | 3 | 1000 | POE | 10 | 5 | 5 | NA | 19 | |
| S5A ($P_{J23106}$-*dnaA* ; nr 7 from left) | 4186 | NA | NA | NA | NA | NA | 3 | 1000 | POE | 10 | 5 | 5 | NA | 19 | |
| S5A ($P_{J23106}$-*dnaA* ; nr 8 from left) | 7495 | NA | NA | NA | NA | NA | 1 | 1000 | POE | 10 | 5 | 5 | NA | 19 | |
| S5A ($P_{J23106}$-*dnaA* ; nr 9 from left) | 8110 | NA | NA | NA | NA | NA | 1 | 1000 | POE | 10 | 5 | 5 | NA | 19 | |
| S5A ($P_{J23106}$-*dnaA* ; nr 10 from left) | 8556 | NA | NA | NA | NA | NA | 3 | 1000 | POE | 10 | 5 | 5 | NA | 19 | |
| S5A ($P_{J23106}$-*dnaA* ; nr 11 from left) | 5471 | NA | NA | NA | NA | NA | 2 | 1000 | POE | 10 | 5 | 5 | NA | 19 | |
| S5A ($P_{J23106}$-*dnaA* ; nr 12 from left) | 8900 | NA | NA | NA | NA | NA | 3 | 1000 | POE | 10 | 5 | 5 | NA | 19 | |
| S5A ($P_{J23106}$-*dnaA* ; nr 13 from left) | 8384 | NA | NA | NA | NA | NA | 1 | 1000 | POE | 10 | 5 | 5 | NA | 19 | |
| S5A ($P_{J23106}$-*dnaA* ; nr 14 from left) | 10198 | NA | NA | NA | NA | NA | 3 | 1000 | POE | 10 | 5 | 5 | NA | 19 | |
| S5A ($P_{J23106}$-*dnaA* ; nr 15 from left) | 7337 | NA | NA | NA | NA | NA | 1 | 1000 | POE | 10 | 5 | 5 | NA | 19 | |
| S5A ($P_{J23106}$-*dnaA* ) | 7983 | NA | NA | NA | NA | NA | 3 | 1000 | POE | 10 | 5 | 5 | NA | 19 | |
| S5A ($P_{wt}$-*dnaA* Δ*DARS1* ) | 6761 | NA | NA | NA | NA | NA | 2 | 1000 | POE | 10 | 5 | 5 | NA | 3 | |
| S5A ($P_{wt}$-*dnaA* Δ*DARS2* ) | 5189 | NA | NA | NA | NA | NA | 1 | 1000 | POE | 10 | 5 | 5 | NA | 2 | |
| S5A ($P_{wt}$-*dnaA* Δ*datA* ) | 6734 | NA | NA | NA | NA | NA | 1 | 1000 | POE | 10 | 5 | 5 | NA | 1 | |
| S5A ($P_{wt}$-*dnaA* Δ*DARS12* Δ*datA* ) | 4474 | NA | NA | NA | NA | NA | 1 | 1000 | POE | 10 | 5 | 5 | NA | 1 | |
| S5A ($P_{wt}$-*dnaA* ) | 8348 | NA | NA | NA | NA | NA | 3 | 1000 | POE | 10 | 5 | 5 | NA | 26 | |
| S5A ($P_{J23106}$-*dnaA* Δ*DARS1* ) | 4209 | NA | NA | NA | NA | NA | 3 | 1000 | POE | 10 | 5 | 5 | NA | 2 | |
| S5A ($P_{J23106}$-*dnaA* Δ*DARS2* ) | 6168 | NA | NA | NA | NA | NA | 1 | 1000 | POE | 10 | 5 | 5 | NA | 2 | |
| S5A ($P_{J23106}$-*dnaA* Δ*datA* ) | 8087 | NA | NA | NA | NA | NA | 1 | 1000 | POE | 10 | 5 | 5 | NA | 2 | |
| S5A ($P_{J23106}$-*dnaA* Δ*DARS12* ) | 6385 | NA | NA | NA | NA | NA | 3 | 1000 | POE | 10 | 5 | 5 | NA | 2 | |
| S5A ($P_{J23106}$-*dnaA* Δ*DARS1* Δ*datA* ) | 4623 | NA | NA | NA | NA | NA | 2 | 1000 | POE | 10 | 5 | 5 | NA | 3 | |
| S5A ($P_{J23106}$-*dnaA* Δ*DARS2* Δ*datA* ) | 8217 | NA | NA | NA | NA | NA | 3 | 1000 | POE | 10 | 5 | 5 | NA | 2 | |
| S5A ($P_{J23106}$-*dnaA* Δ*DARS12* Δ*datA* ) | 7792 | NA | NA | NA | NA | NA | 1 | 1000 | POE | 10 | 5 | 5 | NA | 1 | |
| S5BC ($P_{wt}$-*dnaA* ; nr 1 from left) | 8348 | NA | NA | 3973 | NA | NA | 3 | 1000 | POE | 10 | 5 | 5 | 5 | 26 | |
| S5BC ($P_{wt}$-*dnaA* ; nr 2 from left) | 7759 | NA | NA | 3356 | NA | NA | 2 | 1000 | POE | 10 | 5 | 5 | 5 | 26 | |
| S5BC ($P_{wt}$-*dnaA* ; nr 3 from left) | 6066 | NA | NA | 1950 | NA | NA | 1 | 1000 | POE | 10 | 5 | 5 | 5 | 26 | |
| S5BC ($P_{wt}$-*dnaA* ; nr 4 from left) | 7212 | NA | NA | 2039 | NA | NA | 1 | 1000 | POE | 10 | 5 | 5 | 5 | 26 | |
| S5BC ($P_{wt}$-*dnaA* ; nr 5 from left) | 7924 | NA | NA | 2403 | NA | NA | 1 | 1000 | POE | 10 | 5 | 5 | 5 | 26 | |
| S5BC ($P_{J23106}$-*dnaA* ; nr 6 from left) | 7983 | NA | NA | 5001 | NA | NA | 3 | 1000 | POE | 10 | 5 | 5 | 5 | 19 | |
| S5BC ($P_{J23106}$-*dnaA* ; nr 7 from left) | 4186 | NA | NA | 786 | NA | NA | 3 | 1000 | POE | 10 | 5 | 5 | 5 | 19 | |
| S5BC ($P_{J23106}$-*dnaA* ; nr 8 from left) | 7495 | NA | NA | 1888 | NA | NA | 1 | 1000 | POE | 10 | 5 | 5 | 5 | 19 | |
| S5BC ($P_{J23106}$-*dnaA* ; nr 9 from left) | 8110 | NA | NA | 3963 | NA | NA | 3 | 1000 | POE | 10 | 5 | 5 | 5 | 19 | |
| S5BC ($P_{J23106}$-*dnaA* ; nr 10 from left) | 8556 | NA | NA | 5204 | NA | NA | 3 | 1000 | POE | 10 | 5 | 5 | 5 | 19 | |
| S5BC ($P_{J23106}$-*dnaA* ; nr 11 from left) | 5471 | NA | NA | 1466 | NA | NA | 2 | 1000 | POE | 10 | 5 | 5 | 5 | 19 | |
| S5BC ($P_{J23106}$-*dnaA* ; nr 12 from left) | 8900 | NA | NA | 5607 | NA | NA | 3 | 1000 | POE | 10 | 5 | 5 | 5 | 19 | |
| S5BC ($P_{J23106}$-*dnaA* ; nr 13 from left) | 8384 | NA | NA | 4297 | NA | NA | 1 | 1000 | POE | 10 | 5 | 5 | 5 | 19 | |
| S5BC ($P_{J23106}$-*dnaA* ; nr 14 from left) | 10198 | NA | NA | 6345 | NA | NA | 3 | 1000 | POE | 10 | 5 | 5 | 5 | 19 | |
| S5BC ($P_{J23106}$-*dnaA* ; nr 15 from left) | 7337 | NA | NA | 2171 | NA | NA | 1 | 1000 | POE | 10 | 5 | 5 | 5 | 19 | |
| S5BC ($P_{J23106}$-*dnaA* ) | 7983 | NA | NA | 5001 | NA | NA | 3 | 1000 | POE | 10 | 5 | 5 | 5 | 19 | |
| S5BC ($P_{wt}$-*dnaA* Δ*DARS1* ) | 6761 | NA | NA | 4970 | NA | NA | 2 | 1000 | POE | 10 | 5 | 5 | 5 | 3 | |
| S5BC ($P_{wt}$-*dnaA* Δ*DARS2* ) | 5189 | NA | NA | 2703 | NA | NA | 1 | 1000 | POE | 10 | 5 | 5 | 5 | 2 | |
| S5BC ($P_{wt}$-*dnaA* Δ*datA* ) | 6734 | NA | NA | 2520 | NA | NA | 1 | 1000 | POE | 10 | 5 | 5 | 5 | 1 | |
| S5BC ($P_{wt}$-*dnaA* Δ*DARS12* Δ*datA* ) | 4474 | NA | NA | 2555 | NA | NA | 1 | 1000 | POE | 10 | 5 | 5 | 5 | 1 | |
| S5BC ($P_{wt}$-*dnaA* ) | 8348 | NA | NA | 3973 | NA | NA | 3 | 1000 | POE | 10 | 5 | 5 | 5 | 26 | |
| S5BC ($P_{J23106}$-*dnaA* Δ*DARS1* ) | 4209 | NA | NA | 2210 | NA | NA | 3 | 1000 | POE | 10 | 5 | 5 | 5 | 2 | |
| S5BC ($P_{J23106}$-*dnaA* Δ*DARS2* ) | 6168 | NA | NA | 3039 | NA | NA | 1 | 1000 | POE | 10 | 5 | 5 | 5 | 2 | |
| S5BC ($P_{J23106}$-*dnaA* Δ*datA* ) | 8087 | NA | NA | 2660 | NA | NA | 3 | 1000 | POE | 10 | 5 | 5 | 5 | 2 | |
| S5BC ($P_{J23106}$-*dnaA* Δ*DARS12* ) | 6385 | NA | NA | 3106 | NA | NA | 3 | 1000 | POE | 10 | 5 | 5 | 5 | 2 | |
| S5BC ($P_{J23106}$-*dnaA* Δ*DARS1* Δ*datA* ) | 4623 | NA | NA | 1550 | NA | NA | 2 | 1000 | POE | 10 | 5 | 5 | 5 | 3 | |
| S5BC ($P_{J23106}$-*dnaA* Δ*DARS2* Δ*datA* ) | 8217 | NA | NA | 4981 | NA | NA | 3 | 1000 | POE | 10 | 5 | 5 | 5 | 2 | |
| S5BC ($P_{J23106}$-*dnaA* Δ*DARS12* Δ*datA* ) | 7792 | NA | NA | 4655 | NA | NA | 3 | 1000 | POE | 10 | 5 | 5 | 5 | 1 | |
| S6A (0 ng in wt) | 13773 | NA | NA | NA | NA | NA | 2 | 1000 | Unet | 10 | 5 | 5 | NA | 1 | |
| S6A (0 ng in Δ*DARS12* ) | 7760 | NA | NA | NA | NA | NA | 2 | 1000 | Unet | 10 | 5 | 5 | NA | 1 | |
| S6A (0.5 ng in wt) | 7109 | NA | NA | NA | NA | NA | 2 | 1000 | Unet | 10 | 5 | 5 | NA | 1 | |
| S6A (0.5 ng in Δ*DARS12* ) | 6309 | NA | NA | NA | NA | NA | 2 | 1000 | Unet | 10 | 5 | 5 | NA | 1 | |
| S6A (1 ng in wt) | 6696 | NA | NA | NA | NA | NA | 2 | 1000 | Unet | 10 | 5 | 5 | NA | 3 | |
| S6A (1 ng in Δ*DARS12* ) | 4293 | NA | NA | NA | NA | NA | 2 | 1000 | Unet | 10 | 5 | 5 | NA | 3 | |
| S6A (1 ng in wt) | 5581 | NA | NA | NA | NA | NA | 2 | 1000 | Unet | 10 | 5 | 5 | NA | 3 | |
| S6A (1 ng in Δ*DARS12* ) | 4334 | NA | NA | NA | NA | NA | 2 | 1000 | Unet | 10 | 5 | 5 | NA | 3 | |
| S6A (1 ng in Δ*DARS12* ) | 5942 | NA | NA | NA | NA | NA | 2 | 1000 | Unet | 10 | 5 | 5 | NA | 1 | |
| S6A (1 ng in Δ*DARS12* no repr.) | 7973 | NA | NA | NA | NA | NA | 2 | 1000 | Unet | 10 | 5 | 5 | NA | 1 | |

[a] This is a composite plot consisting of 11078 cells from each of the experiments in Fig. 1D (8 × 11078 = 86624)
