## Supplemental Table S5 for "The coordination of replication initiation with growth rate in *Escherichia coli*"

| Usage | Method | Primer name | Primer 3' -5' | Reference |
|---|---|---|---|---|
| RT-qPCR *dnaA* | qPCR | *dnaA2_f* | GATGAGTTACCAGCCACAGAA | this study |
| RT-qPCR *dnaA* | qPCR | *dnaA2_r* | CGTACCCAATCGAGGACAAA | this study |
| RT-qPCR *dnaA* | qPCR | *hcaT_f* | GCTGCTCGGCTTTCTCATCC | Zhou et al. 2011 |
| RT-qPCR *dnaA* | qPCR | *hcaT_r* | CCAACCACGCTGACCAACC | Zhou et al. 2011 |
| RT-qPCR *dnaA* | qPCR | *cycG_f* | TTGTCGGCGGTGGTGATGTC | Zhou et al. 2011 |
| RT-qPCR *dnaA* | qPCR | *cycG_r* | ATGCGGTGAACTGTGGAATAAACG | Zhou et al. 2011 |
| ori/ter ratios | qPCR | *ori 3_f* | GAGAATATGGCGTACCAGCA | Fernández-Coll et al. 2020 |
| ori/ter ratios | qPCR | *ori 3_r* | AAGACGCAGGTATTTCGCTT | Fernández-Coll et al. 2020 |
| ori/ter ratios | qPCR | *ter 14_f* | GTCATCGGTCAGCGTTAGGT | Fernández-Coll et al. 2020 |
| ori/ter ratios | qPCR | *ter 14_r* | GGTGAATCCTGTCGATGGTC | Fernández-Coll et al. 2020 |
| deletions | λ red DIRex[a] | *cat_midF* | CGACGATTTCCGGCAGTTTC | Näsvall 2017 PLoS One |
| deletions | λ red DIRex[a] | *cat_midR2* | GCCGACATGGAAGCCATCAC | Näsvall 2017 PLoS One |
| deletions | λ red DIRex[a] | *ΔDARS1_DIRex_P1* | GGGAATTGCCCAGGCGGCGGGGGATAGGGGCTGGAGACAG**GAGTTAGAAAACACG**GTGTAGGCTGGAGCTGCTTC | this study |
| deletions | λ red DIRex[a] | *ΔDARS1_DIRex_P2* | AAGCCGCGTATTCTCTCGCTTGCCTCGTGTTTTCTAACTC**CTGTCTCCAGCCCCT**GTGTAGGCTGGAGCTGCTTC | this study |
| deletions | λ red | *ΔDARS2::Acatsac1_f* | TGTGCAGAGTTATAAACAGAGGAAGGGGTGGATAGCCGTT**GTGTAGGCTGGAGCTGCTTC** | this study |
| deletions | λ red | *ΔDARS2::Acatsac1_r* | ATTCAGAATGCTCCGGGTTTTCCCGTAGTAATTCCGTAGA**CATATGAATATCCTCCTTAGTTCC** | this study |
| deletions | λ red | *ΔDARS2_oligo_cure_Acatsac1* | AGGGTTACCGCGTGACGCCTGATTCATCCACAGCTCTCTGTTTACAAGAGTTTGTCAGAGACAACAAAAA | this study |
| deletions | λ red DIRex[a] | *ΔDARS2_DIRex_P1* | GTGCAGAGTTATAAACAGAGGAAGGGGTGGATAGCCGTTT**CTACGGAATTACTAC**GTGTAGGCTGGAGCTGCTTC | this study |
| deletions | λ red DIRex[a] | *ΔDARS2_DIRex_P2* | TATTCAGAATGCTCCGGGTTTTCCCGTAGTAATTCCGTAG**AAACGGCTATCCACC**GTGTAGGCTGGAGCTGCTTC | this study |
| deletions | λ red | *ΔdatA::Acatsac1_f* | GCTGGTTTTTGTTGTCTCTGACAAACTCTTGTAAACAGAG**GTGTAGGCTGGAGCTGCTTC** | this study |
| deletions | λ red | *ΔdatA::Acatsac1_r* | GAAAAAGGGTTACCGCGTGACGCCTGATTCATCCACAGCT**CATATGAATATCCTCCTTAGTTCC** | this study |
| deletions | λ red | *ΔdatA_oligo_cure_Acatsac1* | AGGGTTACCGCGTGACGCCTGATTCATCCACAGCTCTCTGTTTACAAGAGTTTGTCAGAGACAACAAAAA | this study |
| deletions | λ red DUP-in[b] | *Dupin_over_ΔdatA_f* | CTAATGCTACGGCGATATTACGCAGCCAACGCAGGTGACCGT**GTAGGCTGGAGCTGCTTC** | this study |
| deletions | λ red DUP-in[b] | *Dupin_over_ΔdatA_r* | AGACTCGAACTCGCGACCCCGACCTTGGCAAGGTCGTGCT**CATATGAATATCCTCCTTAGTTCC** | this study |
| deletions | λ red | *Δhda::kanR_f* | AGACAAATAATGTACCGACCGGGCAGTGTTGCTGCCCGGT**GCAAGGGCTGCTAAAGGAAG** | this study |
| deletions | λ red | *Δhda::kanR_r* | TCGCGCCGCATCCGACAATAAACACCTTATCTACAACTTC**TTTTGAAGCTGGGGTGGGCG** | this study |
| constitutively expressed *dnaA* | λ red | *Acatsac1_to_galK_f* | AGTCAGCGATATCCATTTTCGCGAATCCGGAGTGTAAGAA**CATATGAATATCCTCCTTAGTTCC** | this study |
| constitutively expressed *dnaA* | λ red | *Acatsac1_to_galK_r* | GACCATCGGGTGCCAGTGCGGGAGTTTCGGTGTAGGCTGGAGCTGCTTCTTCAGCACT**GTGTGTAGGCTGGAGCTGCTTC** | this study |
| constitutively expressed *dnaA* | λ red | *J23106-dnaA_to_galK_f* | AGTCAGCGATATCCATTTTCGCGAATCCGGAGTGTAAGAA**TTTACGGCTAGCTCAGTCCTAGGTATAGTGCTAGC**TCCCGATCGTTTTGCAGGATC | this study |
| constitutively expressed *dnaA* | λ red | *J23100-dnaA_to_galK_f* | AGTCAGCGATATCCATTTTCGCGAATCCGGAGTGTAAGAA**TTGACGGCTAGCTCAGTCCTAGGTACAGTGCTAG**CTCCCGATCGTTTTGCAGGATC | this study |
| constitutively expressed *dnaA* | λ red | *J23101-dnaA_to_galK_f* | TCAGCGATATCCATTTTCGCGAATCCGGAGTGTAAGAAttctagag**TTTACAGCTAGCTCAGTCCTAGGTATAATGCTAGC**tactagagTCCCGATCGTTTTGCAGGATC | this study |
| constitutively expressed *dnaA* | λ red | *J2310X-dnaA_to_galK_save_p22ter_r* | GACCATCGGGTGCCAGTGCGGGAGTTTCGTTCAGCACTGT**GTGTAGGCTGGAGCTGCTTC** | this study |
| constitutively expressed *dnaA* | λ red | *Acatsac1_to_dnaA_f* | TGGTCATTAAATTTTCCAATATGCGGCGTAAATCGTGCCC**CATATGAATATCCTCCTTAGTTCC** | this study |
| constitutively expressed *dnaA* | λ red | *Acatsac1_to_dnaA_r* | TTTGATATCGTGGCTCTCTTCACGCAACTGCTCGATCTTA**GTGTAGGCTGGAGCTGCTTC** | this study |
| constitutively expressed *dnaA* | λ red | *oligo_cure_ΔdnaA::Acatsac1* | ATTAAATTTTCCAATATGCGGCGTAAATCGTGCCC**TTTACGGCTAGCTCAGTCCTAGGTATAGTGCTAGC**TAAGATCGAGCAGTTGCGTGAAGAGAGCCACGATA | this study |
| IPTG inducible DnaA | λ red | *INS_Acatsac1_close_to_plac_F* | GAAGGCGAAGCGGCATGCATTTACGTTGACACCATCGAAT**GTGTAGGCTGGAGCTGCTTC** | this study |
| IPTG inducible DnaA | λ red | *INS_Acatsac1_close_to_plac_R* | CGGCCACCAGAATAGCCTGCGATTCAACCCCTTCTTCGAT**CATATGAATATCCTCCTTAGTTCC** | this study |
| IPTG inducible DnaA | λ red | *synth_part_of_lacZ_dnaA_mcerulean*[c] | ACAGTTGCGCAGCCTGAATGGCGAATGGCGCTTTGCCTGGTTTCCGGCACCAGAAGCGGTGCCGGAAAGCTGGCTGGAGTGCGATCTTCCTGAGGCCGATACTGTCGTC<br>GTCCCCTCAAACTGGCAGATGCACGGTTACGATGCGCCCATCTACACCAACGTGACCTATCCCATTACGGTCAATCCGCCGTTTGTTCCCACGGAGAATCCGACGGGTTG<br>TTACTCGCTCACATTTAATGTTTTAACTAACTAAGTTCGAGTGGAGTCCGCCGTGTCACTTTCGCTTTGGCAGCAGTGTCTTGCCCGATTGCAGGATGAGTTACCAGCCAC<br>AGAATTCAGTATGTGGATACGCCCATTGCAGGCGGAACTGAGCGATAACACGCTGGCCCTGTACGCGCCAAACCGTTTTGTCCTCGATTGGGTACGGGACAAGTACCTTA<br>ATAATATCAATGGACTGCTAACCAGTTTCTGCGGAGCGGATGCCCCACAGCTGCGTTTTGAAGTCGGCACCAAACCGGTGACGCAAACGCCACAATCGGCAGTGACGAG<br>CAACGTCGCGGCCCCTGCACAGGTGGCGCAAACGCAGCCGCAACGTGCTGCGCCTTCTACGCGCTCAGGTTGGGATAACGTCCCGGCCCCGGCAGAACCGACCTATCGT<br>TCTAACGTAAACGTCAAACACACGTTTGATAACTTCGTTGAAGGTAAATCTAACCAACTGGCGCGCGCGGCGGCTCGCCAGGTGGCGGATAACCCTGGCGGTGCCTATAA<br>CCCGTTGTTCCTTTATGGCGGCACGGGTCTGGGTAAAACTCACCTGCTGCATGCGGTGGGTAACGGCATTATGGCGCGCAAGCCGAATGCCAAAGTGGTTTATATGCACT<br>CCGAGCGCTTTGTTCAGGACATGGTTAAAGCCCTGCAAAACAACGCGATCGAAGAGTTTAAACGCTACTACCGTTCCGTAGATGCACTGCTGATCGACGATATTCAGTTT<br>TTTGCTAATAAAGAACGATCTCAGGAAGAGTTTTTCCACACCTTCAACGCCCTGCTGGAAGGTAATCAACAGATCATTCTCACCTCGGATCGCTATCCGAAAGAGATCAA<br>CGGCGTTGAGGATCGTTTGAAATCCCGCTTCGGTTGGGGACTGACTGTGGCGATCGAACCGCCAGAGCTGGAAACCCGTGTGGCGATCCTGATGAAAAAGGCCGACGAA<br>AACGACATTCGTTTGCCGGGCGAAGTGGCGTTCTTTATCGCCAAGCGTCTACGATCTAACGTACGTGAGCTGGAAGGGGCGCTGAACCGCGTCATTGCCAATGCCAACTT<br>TACCGGACGGGCGATCACCATCGACTTCGTGCGTGAGGCGCTGCGCGACTTGCTGGCATTGCAGGAAAAACTGGTCACCATCGACAATATTCAGAAGACGGTGGCGGAG<br>TACTACAAGATCAAAGTCGCGGATCTCCTTTCCAAGCGTCGATCCCGCTCGGTGGCGCGTCCGCGCCAGATGGCGATGGCGCTGGCGAAAGAGCTGACTAACCACAGTCT<br>GCCGGAGATTGGCGATGCGTTTGGTGGCCGTGACCACACGACGGTGCTTCATGCCTGCCGTAAGATCGAGCAGTTGCGTGAAGAGAGCCACGATATCAAAGAAGATTTT<br>TCAAATTTAATCAGAACATTGTCATCGTAAATCTTCCTGGAGAAAATAAGGAGGAAAAAAAAATGAGCAAGGGCGAGGAGCTGTTTACTGGTGTGGTTCCTATCCTGGTC<br>GAACTGGACGGTGATGTCAACGGTCATAAGTTCAGCGTTAGCGGCGAGGGTGAAGGCGACGCAACCTACGGTAAGCTGACCCTGAAGTTCATTTGTACGACGGGTAAGC<br>TGCCGGTGCCTTGGCCTACCCTGGTTACCACCCTGACCTGGGGTGTTCAATGCTTCGCTCGCTATCCAGACCACATGAAGCAGCACGATTTTTTTAAGAGCGCCATGCCAG<br>AGGGTTACGTTCAGGAGCGCACCATTTTCTTTAAAGATGATGGTAATTACAAGACTCGCGCAGAGGTGAAGTTTGAAGGCGACACTCTGGTGAATCGTATTGAGCTGAA<br>GGGCATCGATTTTAAGGAAGATGGTAATATCCTGGGCCATAAACTGGAATATAATGCGATTAGCGATAATGTGTATATCACCGCCGATAAACAAAAGAATGGTATTAAG | this study |
| IPTG inducible DnaA | λ red | *Acatsac1_mutation_fix_dnaA_f* | CGTCGGCCTTTTTCATCAGGATCGCCACACGGGTTTCCAG**CATATGAATATCCTCCTTAGTTCC** | this study |
| IPTG inducible DnaA | λ red | *Acatsac1_mutation_fix_dnaA_r* | GGCCACCAAACGCATCGCCAATCTCCGGCAGACTGTGGTT**CATATGAATATCCTCCTTAGTTCC** | this study |
| IPTG inducible DnaA | λ red | *exchange_dnaA::Acatsac1_for_dnaA_f* | ATATCGTGGCTCTCTTCAC | this study |
| IPTG inducible DnaA | λ red | *exchange_dnaA::Acatsac1_for_dnaA_r* | CGTGTCACTTTCGCTTTG | this study |
| *parS/parB* terminus tag | λ red | *parS-cat_to_ydbL_f* | GAGCTGCTTGAAACTCGTAGCGATCTTTTCTGAGGTGATG**TTTTCACCACGCCAATTTCA** | this study |
| *parS/parB* terminus tag | λ red | *parS-cat_to_ydbL_r* | CGCCTGTAGCGCAGCTTCACTTTCTGTTTCAGAAAGTGCC**GTGTAGGCTGGAGCTGCTTC** | this study |

[a] Näsvall 2017 PLoS One

[b] Näsvall, Knöppel, Andersson 2017 NAR

[c] the synthesis caused the dnaA(G265T; Ala89Ser) mutation. The primers served to fix this.
