## Supplemental Figures and Tables for "The coordination of replication initiation with growth rate in *Escherichia coli*"

Figures S1 – S7

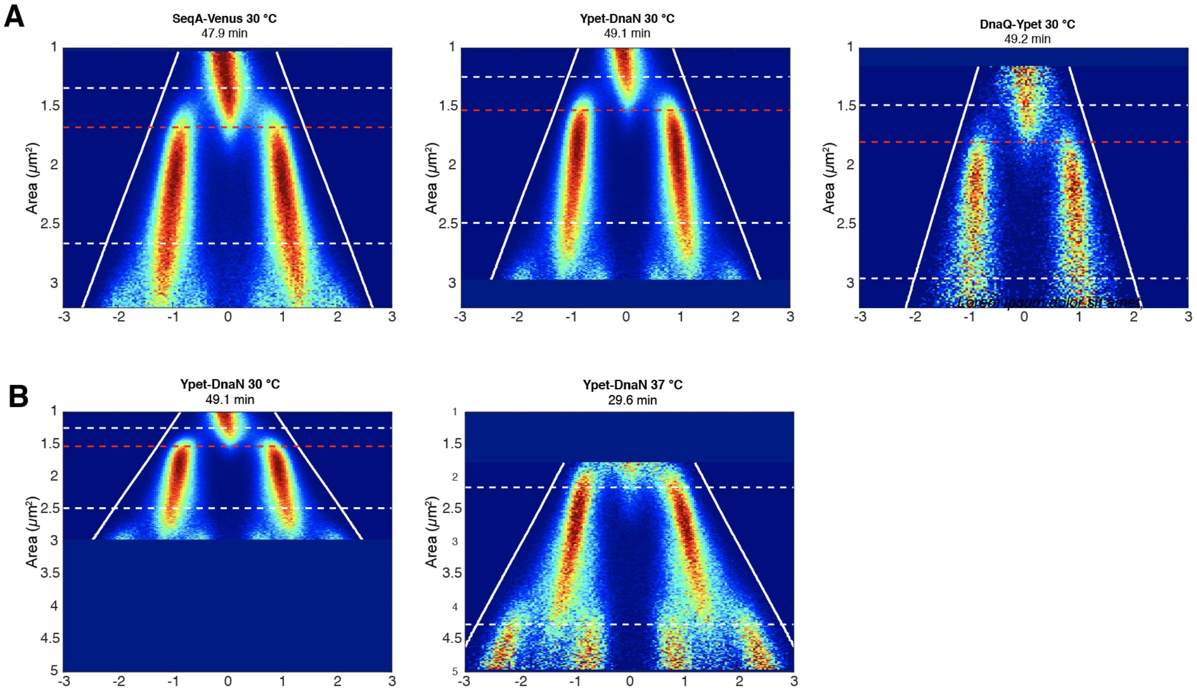


***Figure S1. Comparison of different replication tracking markers and the influence of temperature on replication. (A)*** *Fork plots for the three different markers of the replication fork.* ***(B)*** *The strain carrying YPet-DnaN and parS-ter/ParB-mCherry was grown at 30 and 37 °C. All cells were grown in M9 0.4% succinate medium supplemented with amino acids.*

## A

| | $P_{wt}$-*dnaA* | $P_{J23106}$-*dnaA* | $P_{J23100}$-*dnaA* | $P_{J23101b}$-*dnaA* |
|---|---|---|---|---|
| *oriC* content[a] | 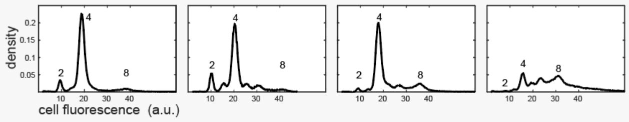 | | | |
| Doubling time[b] | 44.6 (0.04) | 45.6 (0.03) | 44.7 (0.01) | 46.3 (na) |
| [*dnaA* mRNA][c] | 1.00 (0.19) | 1.03 (0.45) | 2.31 (na) | 2.67 (0.29) |
| [DnaA protein][d] | 1.00 (na) | 0.99 (na) | 2.04 (na) | 2.66 (na) |

## B

Fork plot
Ref. strain

Fork plot
Mutant strain

## C

Supercell init. area
Ref. strain

Supercell init. area
Mutant strain

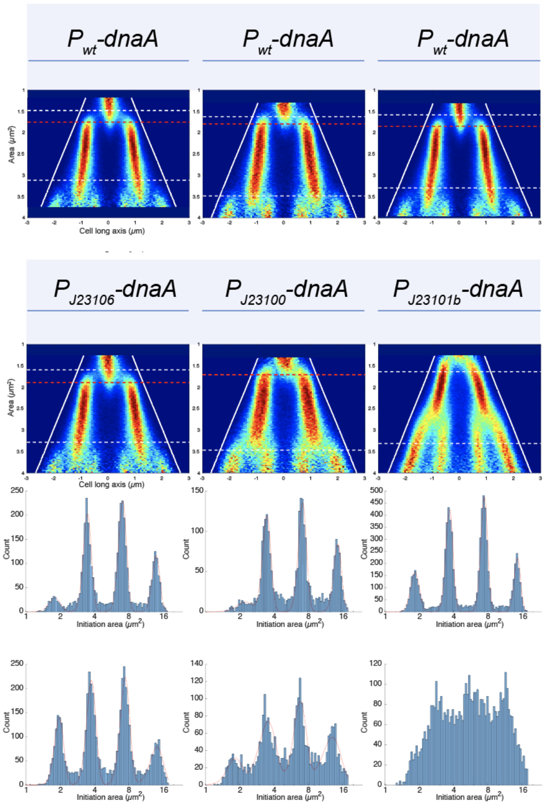


## D

| | | | |
|---|---|---|---|
| Init. area ref. | 1.75 (0.12) | 1.79 (0.11) | 1.85 (0.10) |
| Init. area mut. | 1.89 (0.13) | 1.71 (0.18) | na (na) |
| Birth area ref. | 1.48 (0.18) | 1.62 (0.17) | 1.58 (0.17) |
| Birth area mut. | 1.59 (0.20) | 1.71 (0.21) | 1.63 (0.23) |
| Division area ref. | 3.12 (0.15) | 3.49 (0.15) | 3.30 (0.15) |
| Division area mut. | 3.28 (0.17) | 3.47 (0.17) | 3.31 (0.19) |
| Doubling time ref. | 46 (0.10) | 44 (0.07) | 45 (0.07) |
| Doubling time mut. | 46 (0.09) | 47 (0.08) | 46 (0.10) |

***Figure S2*. *Comparative measurements of strains carrying $P_{wt}$-dnaA, $P_{J23106}$-dnaA, $P_{J23100}$-dnaA and $P_{J23101b}$-dnaA.*** *(**A**) [a]Results of rif-runout experiments. [b]Bioscreen plate reader (n = 2 or 3), [c]RT-qPCR (n = 2–6), and [d]LC-MS/MS (n = 2). n is the number of biological replicates. The strains used for rifampicin run-out experiments carried no other modifications on the chromosome, whereas all other data were generated from strains additionally carrying the chromosomal cobA::seqA-venus construct. (**B–D**) Microfluidic measurements (**B**) The same fork plots as displayed in Figure 2A, but with the addition of the reference strain that was run in the same experiment (top). (**C**) Distributions of initiation areas from super-cells fitted with a sum of four Gaussians. It was not possible to fit Gaussians to the distribution of the $P_{J23101b}$ strain. (**D**) Average initiation areas based on fits in (C) and the average birth areas, division areas and doubling times. Coefficients of variation for the size and time distributions of a single experiment are displayed within the parentheses. For all experiments, the cells were grown in M9 0.4% succinate supplemented with amino acids at 30 °C.*

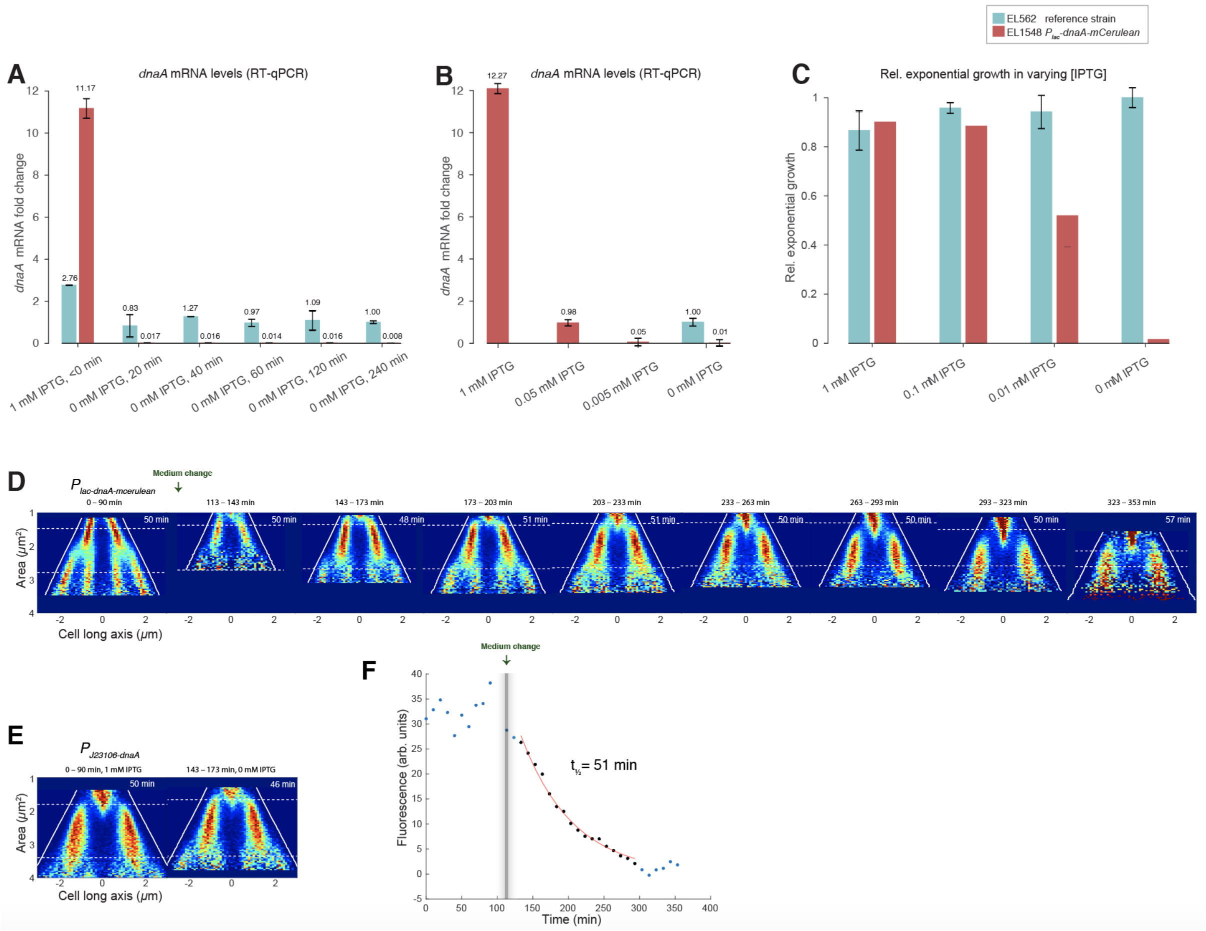


***Figure S3*. *Measurements on the $P_{lac}$-dnaA-mCerulean strain*.** *(**A)** RT-qPCR. Relative dnaA mRNA levels at varying time points before and after medium swap (30 °C M9 0.4% succinate medium supplemented with amino acids, +IPTG to -IPTG). All values are related to EL562 at 240 min after swap. Reported values represent the mean (±SD) of three (EL1548) or two (EL562) independent experiments. (**B)** RT-qPCR. Relative dnaA mRNA levels at varying IPTG concentrations. EL1548 at 0 mM IPTG was taken 240 min after a medium swap from 1 mM IPTG. All values are related to EL562 grown in 0 mM IPTG. Reported values represent the mean (±SD) of four independent replicates. (**C)** Relative exponential growth rate in succinate medium at 37 °C. EL562 grown in 0 mM IPTG serves as control. Reported values represent the mean (±SD) of two (EL1548) or three (EL562) independent replicates. (**D–F)** Experimental repeat of Figures 3A, 3B and 3E.*

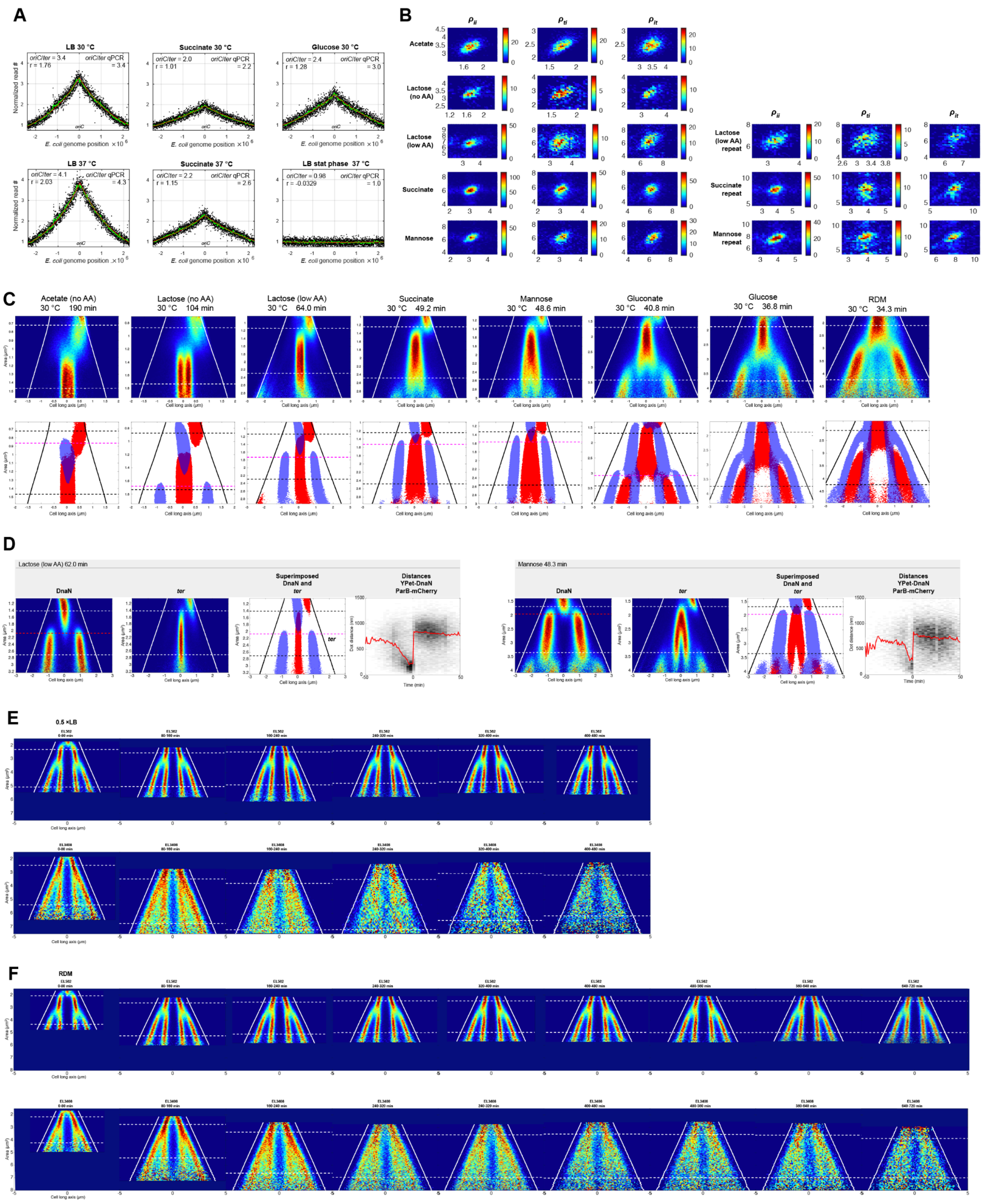


***Figure S4*. Marker frequency analysis, termination-initiation correlations, replisome-ter distances and Δhda medium swap experiments.** *(**A**) Marker frequency analysis. Normalized number of mapped reads (y-axis) for each chromosome region (x-axis) for a wt strain grown at different conditions. The genome coordinate is centered around oriC. The number of reads was binned into regions of either ~1.5 kb (black dots) or ~30 kb (green line) and normalized to the leftmost bin in each case. Red lines are the results of regressions of the 30 kb binned data (green line) to the function $y = B * 2^{(-C*x/\tau)}$. Here, $x$ is a normalized genome coordinate where $x = 0$ at oriC and $x = 1$ on the opposite side. Inset in each panel: oriC/ter ratios are calculated from regression as $2^{(C/\tau)}$. $r =$ C-period / generation time. qPCR determinations of oriC/ter ratios are shown to the left in each panel. Reported values are the mean of the values from two oriC PCR replicates divided by the same ter PCR. (**B**) Matched initiation and termination events from super-cells clustered for all growth conditions where correlations between replication events were determined. The colorbar shows the number of*

*matched events in each bin. (**C**) Fork plots of a strain that simultaneously carries fluorescent markers for replication (a YPet-DnaN translational fusion; Figure 1C) and terminus (the ParB-mCherry/parS system, where parS is placed in the terminus region; first row). In the second row, the results from YPet-DnaN (Figure 1C) have been superimposed onto the results from the terminus marker. The results are displayed as filled contours. Lines as in Figure 1C, except white lines are now black and the red line is changed to magenta. (**D**) Distances between replisomes and termini in single cells grown in either M9 10 mM lactose with low amounts of amino acids or M9 20 mM mannose supplemented with amino acids. First three columns: Fork plots of a strain that carries fluorescent markers for replication (a YPet-DnaN translational fusion; first column) and terminus (the ParB-mCherry/parS system, where parS is placed in the terminus region; second column). In the third column, filled contours of the two fork plots have been superimposed. The lines are the same as in (C). Fourth column: Heat-map of 2D histograms for distances between YPet-DnaN and ParB-mCherry foci in single cells and relative time from YPet-DnaN track disappearance. Red lines indicate the median distance over time. (**E**) Fork plots for wt and Δhda for the full experimental duration after swapping from M9 0.4% succinate medium supplemented with amino acids to 0.5× LB medium. (**F**) Same as (E) but the swap was to RDM.*

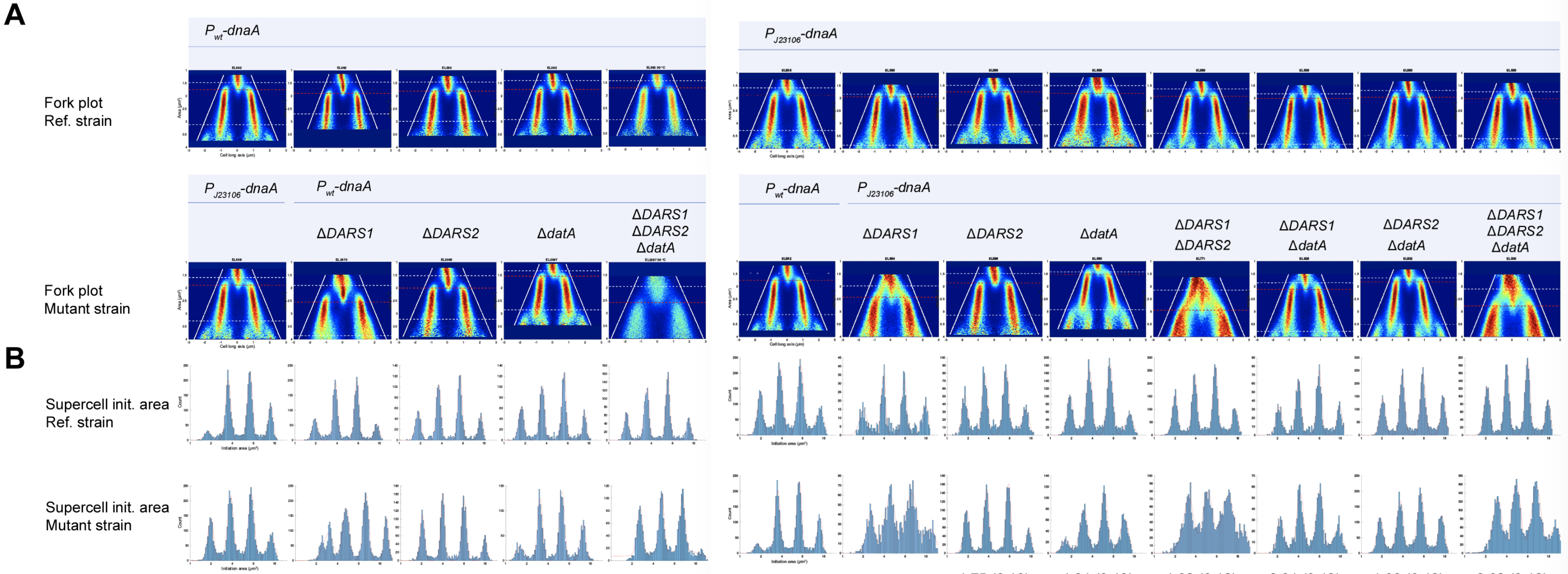


**C**

| | $P_{J23106}$-dnaA | ΔDARS1 | ΔDARS2 | ΔdatA | ΔDARS1 ΔDARS2 ΔdatA |
|---|---|---|---|---|---|
| Init. area ref. | 1.75 (0.12) | 1.90 (0.11) | 1.81 (0.10) | 1.74 (0.11) | 1.73 (0.11) |
| Init. area mut. | 1.89 (0.13) | 2.55 (0.13) | 2.00 (0.09) | 1.54 (0.11) | 2.63 (0.11) |
| Birth area ref. | 1.48(0.18) | 1.46 (0.18) | 1.46 (0.15) | 1.40 (0.18) | 1.42 (0.17) |
| Birth area mut. | 1.59 (0.20) | 1.90 (0.21) | 1.52 (0.18) | 1.33 (0.18) | 1.95 (0.19) |
| Division area ref. | 3.12 (0.15) | 2.70 (0.16) | 2.99 (0.14) | 2.92 (0.16) | 2.95 (0.16) |
| Division area mut. | 3.28 (0.17) | 3.86 (0.16) | 3.22 (0.16) | 2.85 (0.16) | 4.07(0.18) |
| Doubling time ref. | 46 (0.10) | 51 (0.10) | 47 (0.07) | 44 (0.08) | 45 (0.07) |
| Doubling time mut. | 46 (0.09) | 46 (0.10) | 45 (0.08) | 44 (0.08) | 44 (0.07) |

| $P_{wt}$-dnaA | ΔDARS1 | ΔDARS2 | ΔdatA | ΔDARS1 ΔDARS2 | ΔDARS1 ΔdatA | ΔDARS2 ΔdatA | ΔDARS1 ΔDARS2 ΔdatA |
|---|---|---|---|---|---|---|---|
| 1.89 (0.13) | 1.95 (0.10) | 1.75 (0.13) | 1.81 (0.13) | 1.92 (0.12) | 2.01 (0.12) | 1.96 (0.12) | 2.02 (0.12) |
| 1.75 (0.12) | 2.43 (0.24) | 1.86 (0.11) | 1.52 (0.18) | 2.94 (0.23) | 2.12 (0.16) | 1.82 (0.15) | 2.78 (0.21) |
| 1.59 (0.18) | 1.84 (0.20) | 1.49 (0.20) | 1.50 (0.21) | 1.63 (0.19) | 1.83 (0.18) | 1.69 (0.20) | 1.75 (0.19) |
| 1.48 (0.18) | 1.98 (0.21) | 1.49 (0.18) | 1.42 (0.21) | 2.15 (0.20) | 1.86 (0.18) | 1.66 (0.19) | 2.10 (0.21) |
| 3.28 (0.17) | 3.87 (0.17) | 3.06 (0.17) | 3.05 (0.19) | 3.40 (0.16) | 3.83 (0.15) | 3.48 (0.17) | 3.61 (0.16) |
| 3.12 (0.15) | 4.07 (0.18) | 3.14 (0.17) | 2.92 (0.19) | 4.35 (0.18) | 3.79 (0.15) | 3.50 (0.17) | 4.32 (0.18) |
| 46 (0.09) | 49 (0.07) | 45 (0.08) | 45 (0.09) | 46 (0.07) | 44 (0.06) | 50 (0.07) | 47 (0.06) |
| 46 (0.10) | 48 (0.07) | 44 (0.08) | 46 (0.10) | 47 (0.07) | 46 (0.07) | 49 (0.07) | 47 (0.08) |

**D**

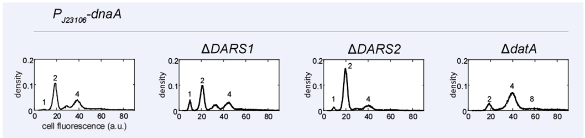

***Figure S5. Comparison of different DnaA-ATP/ADP regulatory mutants with wild-type expression of DnaA**. (**A–C**) Microfluidic experiments. (**A)** Fork plots of different mutants constructed in $P_{wt}$-dnaA background (left section) and $P_{J23106}$-dnaA background (right section; same as Figure 5A). As shown in Figure 5A with the addition of the reference strain that was included in the same experiment is shown on the top row. (**B**) Same as Figure S2B but for strains in (A). (**C**) Same as Figure S2C, but for strains in (A). (**D**) Results of rif-runout experiments on oriC content. In all experiments cells were grown in M9 0.4% succinate medium supplemented with amino acids at 30 °C.*

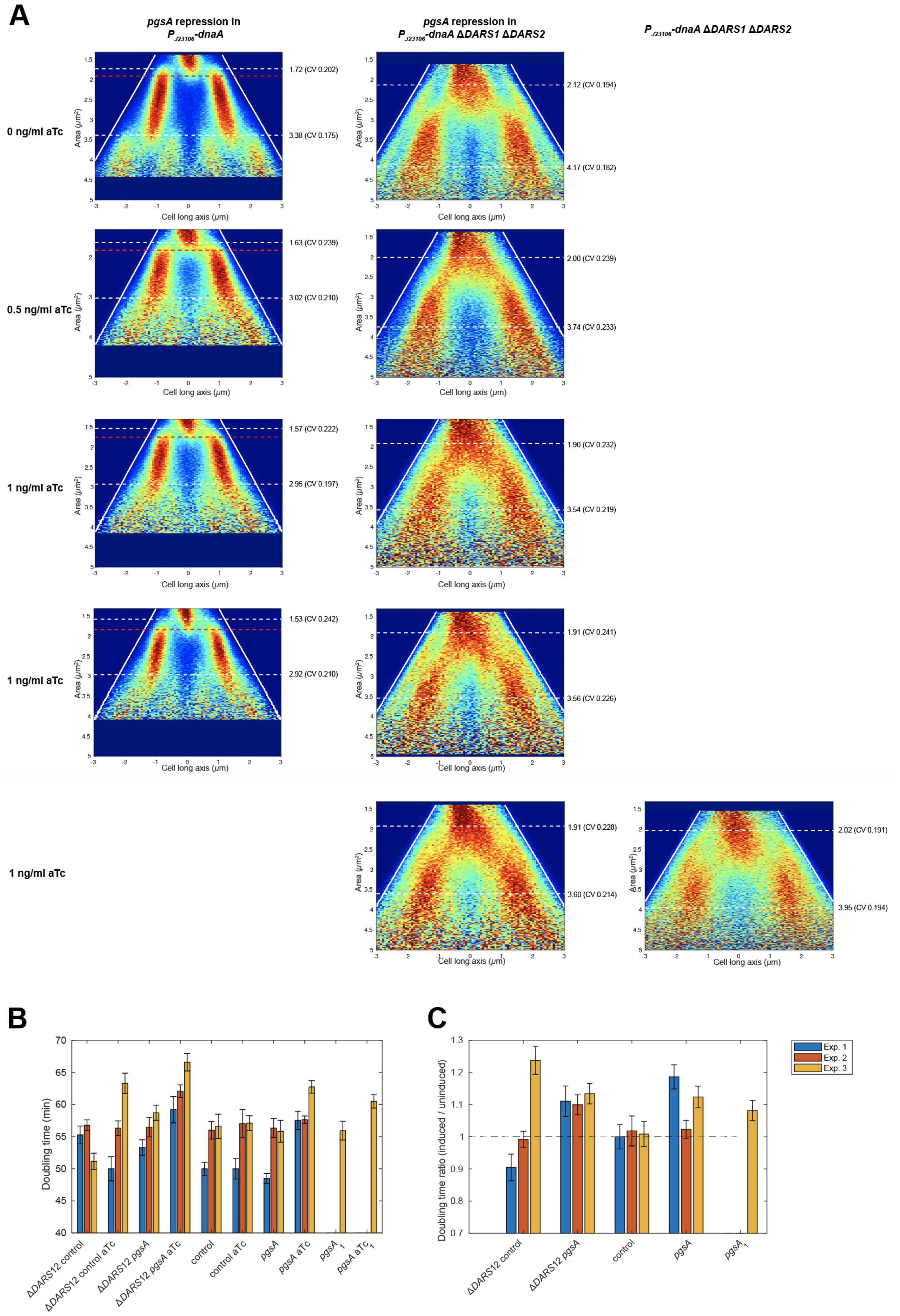
A
pgsA repression in P J23106-dnaA
pgsA repression in P J23106-dnaA ΔDARS1 ΔDARS2
P J23106-dnaA ΔDARS1 ΔDARS2
0 ng/ml aTc
0.5 ng/ml aTc
1 ng/ml aTc
1 ng/ml aTc
1 ng/ml aTc
1.72 (CV 0.202)
3.38 (CV 0.175)
2.12 (CV 0.194)
4.17 (CV 0.182)
1.63 (CV 0.239)
3.02 (CV 0.210)
2.00 (CV 0.239)
3.74 (CV 0.233)
1.57 (CV 0.222)
2.95 (CV 0.197)
1.90 (CV 0.232)
3.54 (CV 0.219)
1.53 (CV 0.242)
2.92 (CV 0.210)
1.91 (CV 0.241)
3.56 (CV 0.226)
1.91 (CV 0.228)
3.60 (CV 0.214)
2.02 (CV 0.191)
3.95 (CV 0.194)
Area (μm²)
Cell long axis (μm)
B
Doubling time (min)
ΔDARS12 control
ΔDARS12 control aTc
ΔDARS12 pgsA
ΔDARS12 pgsA aTc
control
control aTc
pgsA
pgsA aTc
pgsA 1
pgsA aTc 1
C
Doubling time ratio (induced / uninduced)
ΔDARS12 control
ΔDARS12 pgsA
control
pgsA
pgsA 1
Exp. 1
Exp. 2
Exp. 3

***Figure S6*. *pgsA repression in the presence and absence of DARS1 and DARS2.*** *(**A**) Fork plots for all replicates of pgsA repression with different levels of anhydrotetracycline (aTc), with and without DARS1 and DARS2 as well as a control with ΔDARS1 ΔDARS2 without pgsA repression. (**B**) Experiment where the doubling times of the strains with and without DARS1 and DARS2 were determined in E-flasks with and without pgsA repression. The experiment was performed in M9 0.4% succinate medium supplemented with amino acids at 30 °C. (**C**) Uninduced-induced doubling time ratios for the same strains as in (B).*

### Tables S1 – S6

***Table S1***. *Media.*

| Medium | Supplement | Abbreviation | Figure |
|---|---|---|---|
| M9 minimal medium | 0.4% succinate and 1x RPMI 1640[b] | Succinate | standard medium[c] |
| M9 minimal medium | 0.4% acetate | Acetate (no AA) | 1CD, 4CD, S4BC |
| M9 minimal medium | 10 mM lactose | Lactose (no AA) | 1CD, 4D, S4BC |
| M9 minimal medium | 10 mM lactose and 0.025x RPMI 1640[b] | Lactose (low AA) | 1CD, 4D, S4BCD |
| M9 minimal medium | 20 mM mannose and 1x RPMI 1640[b] | Mannose | 1CD, 4D, S4BCD |
| M9 minimal medium | 20 mM gluconate and 1x RPMI 1640[b] | Gluconate | 1CD, S4C |
| M9 minimal medium | 0.4% glucose and 1x RPMI 1640[b] | Glucose | 1CD, S4C |
| RDM | - | RDM | 1CD , 4AB, S4F |
| LB | - | LB | S4A |
| 0.5× LB | Sigma $H_2O$ | 0.5× LB | 4AB, S4E |

[a] We additionally supplemented all media used in microscopy experiments with the surfactant Gibco® Pluronic® F-108. 0.18× (51 μg/ml) for RDM and 0.5× (153 μg/ml) LB, and 0.06× for all other media.

[b] RPMI 1640 Amino Acids Solution (50×; R7131 Sigma-Aldrich).

[c] Used in experiments in most figures.

***Table S2***. *Correlations between initiation, termination, division and growth rate for cells in Figure 1C. $\rho_{II}$, $\rho_{IT}$, $\rho_{TI}$, and $\rho_{TI,model}$ are correlation coefficients between cell sizes at two subsequent initiations ($\rho_{II}$), initiation and the corresponding termination ($\rho_{TI}$) and termination and the next-coming initiation ($\rho_{II}$). $\rho_{TI,model}$ is a predicted correlation between termination and initiation based on the experimentally observed $\rho_{II}$ and $\rho_{IT}$ (see main text for description). $\rho_{BI}$ is the correlation coefficient between cell sizes at birth and the subsequent initiation, $\rho_{ID}$ for initiation and the subsequent division and $\rho_{TD}$ for termination and the subsequent division. $\rho_{DD}$ is the correlation coefficient between cell sizes at two subsequent divisions. $\rho_{GG}$ is the correlation coefficient between growth rates in two consecutive generations.*

| Medium | $\rho_{II}$ | $\rho_{IT}$ | $\rho_{TI}$ | $\rho_{TI,\ model}$ | $\rho_{BI}$ | $\rho_{ID}$ | $\rho_{TD}$ | $\rho_{DD}$ | $\rho_{GG}$ |
|---|---|---|---|---|---|---|---|---|---|
| Acetate (no AA) | 0.51 | 0.64 | 0.31 | 0.33 | 0.27 | 0.44 | 0.48 | 0.37 | 0.48 |
| Lactose (no AA) | 0.54 | 0.54 | 0.65 | 0.29 | 0.24 | 0.55 | 0.55 | 0.52 | 0.19 |
| Lactose (low AA) | 0.43 | 0.27 | 0.53 | 0.11 | 0.08 | 0.19 | - | 0.41 | 0.15 |
| Succinate | 0.36 | 0.33 | 0.24 | 0.12 | 0.03 | 0 | 0.06 | 0.56 | 0.35 |
| Mannose | 0.51 | 0.5 | 0.55 | 0.26 | 0.16 | 0 | 0.09 | 0.52 | 0.28 |
| Lactose (low AA)[a] | 0.47 | 0.31 | 0.52 | 0.15 | 0.08 | 0.25 | 0.55 | 0.37 | 0.21 |
| Succinate[a] | 0.34 | 0.093 | 0.46 | 0.035 | 0 | 0 | 0 | 0.49 | 0.38 |
| Mannose[a] | 0.56 | 0.45 | 0.64 | 0.26 | 0 | 0.04 | 0.03 | 0.54 | 0.22 |

[a]repeat experiment

***Table S3***. *Strain list.*

| Strain number | Strain | Genotype | Ref |
|---|---|---|---|
| EL544 (wt) | wt | *E. co* MG1655 BW25993 Δ*phi80 rph+* | This study |
| **Derivatives of EL544** | | | |
| EL421 | *J23106-dnaA* Δ*dnaA* | Δ*galK* ::*J23106-dnaA* Δ*pdnaA-dnaA* ::*J23106-dnaA* (123 nt) | This study |
| EL838 | *J23100-dnaA* Δ*dnaA* | Δ*galK* ::*J23100-dnaA* Δ*pdnaA-dnaA* ::*J23106-dnaA* (123 nt) | This study |
| EL787[a] | *J23101b-dnaA* Δ*dnaA* | Δ*galK* ::*J23101b-dnaA* Δ*pdnaA-dnaA* ::*J23106-dnaA* (123 nt) | This study |
| EL840 | *J23106-dnaA* Δ*dnaA* Δ*DARS1* | Δ*galK* ::*J23106-dnaA* Δ*pdnaA-dnaA* ::*J23106-dnaA* (123 nt) Δ*DARS1* | This study |
| EL548 | *J23106-dnaA* Δ*dnaA* Δ*DARS2* | Δ*galK* ::*J23106-dnaA* Δ*pdnaA-dnaA* ::*J23106-dnaA* (123 nt) Δ*DARS2* | This study |
| EL547 | *J23106-dnaA* Δ*dnaA* Δ*datA* | Δ*galK* ::*J23106-dnaA* Δ*pdnaA-dnaA* ::*J23106-dnaA* (123 nt) Δ*datA* | This study |
| EL562 | *seqA-venus* | Δ*cobA* ::*seqA-venus-FRTcatFRT* | This study |
| EL2470 | *seqA-venus* Δ*DARS1* | Δ*cobA::seqA-venus-FRTcatFRT* Δ*DARS1* | This study |
| EL2095 | *seqA-venus* Δ*DARS2* | Δ*cobA::seqA-venus-FRTcatFRT* Δ*DARS2* | This study |
| EL2097 | *seqA-venus* Δ*datA* | Δ*cobA::seqA-venus-FRTcatFRT* Δ*datA* | This study |
| EL2297 | *seqA-venus* Δ*DARS1* Δ*DARS2* Δ*datA* | Δ*cobA::seqA-venus-FRTcatFRT* Δ*DARS1* Δ*DARS2* Δ*datA* | This study |
| EL862 | *seqA-venus J23100-dnaA* Δ*dnaA* | Δ*cobA* ::*seqA-venus-FRTcatFRT* Δ*galK* ::*J23100-dnaA* Δ*pdnaA-dnaA* ::*J23106-dnaA* (123 nt) | This study |
| EL868[a] | *seqA-venus J23101b-dnaA* Δ*dnaA* | Δ*cobA* ::*seqA-venus-FRTcatFRT* Δ*galK* ::*J23101b-dnaA* Δ*pdnaA-dnaA* ::*J23106-dnaA* (123 nt) | This study |
| EL558 | *seqA-venus J23106-dnaA* Δ*dnaA* | Δ*cobA* ::*seqA-venus-FRTcatFRT* Δ*galK* ::*J23106-dnaA* Δ*pdnaA-dnaA* ::*J23106-dnaA* (123 nt) | This study |
| EL864 | *seqA-venus J23106-dnaA* Δ*dnaA* Δ*DARS1* | Δ*cobA* ::*seqA-venus-FRTcatFRT* Δ*galK* ::*J23106-dnaA* Δ*pdnaA-dnaA* ::*J23106-dnaA* (123 nt) Δ*DARS1* | This study |
| EL586 | *seqA-venus J23106-dnaA* Δ*dnaA* Δ*DARS2* | Δ*cobA* ::*seqA-venus-FRTcatFRT* Δ*galK* ::*J23106-dnaA* Δ*pdnaA-dnaA* ::*J23106-dnaA* (123 nt) Δ*DARS2* | This study |
| EL560 | *seqA-venus J23106-dnaA* Δ*dnaA* Δ*datA* | Δ*cobA* ::*seqA-venus-FRTcatFRT* Δ*galK* ::*J23106-dnaA* Δ*pdnaA-dnaA* ::*J23106-dnaA* (123 nt) Δ*datA* | This study |
| EL771 | *seqA-venus J23106-dnaA* Δ*dnaA* Δ*DARS1* Δ*DARS2* | Δ*cobA* ::*seqA-venus-FRTcatFRT* Δ*galK* ::*J23106-dnaA* Δ*pdnaA-dnaA* ::*J23106-dnaA* (123 nt) Δ*DARS1* Δ*DARS2* | This study |
| EL929 | *seqA-venus J23106-dnaA* Δ*dnaA* Δ*DARS1* Δ*datA* | Δ*cobA* ::*seqA-venus-FRTcatFRT* Δ*galK* ::*J23106-dnaA* Δ*pdnaA-dnaA* ::*J23106-dnaA* (123 nt) Δ*DARS1* Δ*datA* | This study |
| EL502 | *seqA-venus J23106-dnaA* Δ*dnaA* Δ*DARS2* Δ*datA* | Δ*cobA* ::*seqA-venus-FRTcatFRT* Δ*galK* ::*J23106-dnaA* Δ*pdnaA-dnaA* ::*J23106-dnaA* (123 nt) Δ*DARS2* Δ*datA* | This study |
| EL500 | *seqA-venus J23106-dnaA* Δ*dnaA* Δ*DARS1* Δ*DARS2* Δ*datA* | Δ*cobA* ::*seqA-venus-FRTcatFRT* Δ*galK* ::*J23106-dnaA* Δ*pdnaA-dnaA* ::*J23106-dnaA* (123 nt) Δ*DARS1* Δ*DARS2* Δ*datA* | This study |
| EL1548 | *seqA-venus plac-dnaA-mcerulean* Δ*dnaA* | Δ*cobA* ::*seqA-venus-FRTcatFRT* Δ*lacZYA* ::*dnaA-mcerulean3-FRT* Δ*pdnaA-dnaA* ::*J23106-dnaA* (123 nt) | This study |
| EL2822 | *seqA-venus plac-dnaA-mcerulean* Δ*dnaA* Δ*DARS1* Δ*DARS2* Δ*datA* | Δ*cobA* ::*seqA-venus-FRTcatFRT* Δ*lacZYA* ::*dnaA-mcerulean3-FRT-kanR-FRT* Δ*pdnaA-dnaA* ::*J23106-dnaA* (123 nt) Δ*DARS1* Δ*DARS2* Δ*datA* | This study |
| EL3242 | *seqA-venus J23106-dnaA* Δ*dnaA tetR-dCas9 /pguide-control* | Δ*cobA* ::*seqA-venus-FRT* Δ*galK* ::*J23106-dnaA* Δ*pdnaA-dnaA* ::*J23106-dnaA* (123 nt) *tetR-PRoger-PLtetO1-dcas9 /pGuide7-WTO1a* | This study |
| EL3244 | *seqA-venus J23106-dnaA* Δ*dnaA tetR-dCas9 /pguide-pgsA* | Δ*cobA* ::*seqA-venus-FRT* Δ*galK* ::*J23106-dnaA* Δ*pdnaA-dnaA* ::*J23106-dnaA* (123 nt) *tetR-PRoger-PLtetO1-dcas9 /pGuide7-pgsA* | This study |
| EL3298 | *seqA-venus J23106-dnaA* Δ*dnaA* Δ*DARS1* Δ*DARS2 tetR-dCas9 /pguide- contrc* | Δ*cobA* ::*seqA-venus-FRT* Δ*galK* ::*J23106-dnaA* Δ*pdnaA-dnaA* ::*J23106-dnaA* (123 nt) *tetR-PRoger-PLtetO1-dcas9* Δ*DARS1* Δ*DARS2 /pGuide7-WTC* | This study |
| EL3299 | *seqA-venus J23106-dnaA* Δ*dnaA* Δ*DARS1* Δ*DARS2 tetR-dCas9 /pguide-pgsA* | Δ*cobA* ::*seqA-venus-FRT* Δ*galK* ::*J23106-dnaA* Δ*pdnaA-dnaA* ::*J23106-dnaA* (123 nt) *tetR-PRoger-PLtetO1-dcas9* Δ*DARS1* Δ*DARS2 /pGuide7-pgsA* | This study |
| EL3308 | *seqA-venus J23106-dnaA* Δ*dnaA* Δ*DARS1* Δ*DARS2 J23106-pgsA* | Δ*cobA* ::*seqA-venus-FRTcatFRT* Δ*galK* ::*J23106-dnaA* Δ*pdnaA-dnaA* ::*J23106-dnaA* (123 nt) Δ*DARS1* Δ*DARS2* Δ*intS* ::*J23106-pgsA* | This study |
| EL3408 | *seqA-venus* Δ*hda* | Δ*cobA* ::*seqA-venus-FRT lac*$^{-}$ Δ*hda* ::*kanR* | This study |
| EL2931 | *seqA-venus* Δ*ydbL* ::*parS* Δ*gtrA* ::*mCherry-parB* | Δ*cobA* ::*seqA-venus-FRTcatFRT* Δ*ydbL* ::*parS-FRT-cat-FRT* Δ*gtrA* ::*p58-mCherry-parB-SpR* | This study |
| EL2990 | *ypet-dnaN* Δ*ydbL* ::*parS* Δ*gtrA* ::*mCherry-parB* | *kan-ypet-dnaN* Δ*ydbL* ::*parS-FRT-cat-FRT* Δ*gtrA* ::*p58-mCherry-parB-SpR* | This study |
| EL2938 | *dnaQ-ypet* Δ*ydbL* ::*parS* Δ*gtrA* ::*mCherry-parB* | *dnaQ-yfp-FRT-kanR-FRT* Δ*ydbL* ::*parS-FRT-cat-FRT* Δ*gtrA* ::*p58-mCherry-parB-SpR* | This study |

[a]The J23101b promoter is based on J23101 but carries one missmatch (TTTACAGCTAGCTCAGTCCTAGGTATA**A**TGCTAGC instead of TTTACAGCTAGCTCAGTCCTAGGTAT**T**ATGCTAGC

***Table S4***. *Number of cells (or equivalent) that were used to create the plots in the main text and supplement. Microscopy configuration numbers, chip size, segmentation algorithm, cell selection criteria and the number of repeats are also listed. The table is only available as a separate .pdf file.*

***Table S5***. *Primers and other synthesized DNA. The table is only available as a separate .pdf file.*

***Table S6***. *Quantification of relative DnaA protein levels through TMT LC-MS/MS.*

| Strain | Genotype | Condition | Sample A | Sample B |
|---|---|---|---|---|
| EL562 | ref | succinate 30 °C | 1.000[c] | |
| EL1548 | *p* $_{lac}$*-dnaA* | succinate 30 °C 1 mM IPTG | 7.381 | 4.686 |
| EL1548 | *p* $_{lac}$*-dnaA* | succinate 30 °C 75 µM IPTG | 2.348 | 2.239 |
| EL1548 | *p* $_{lac}$*-dnaA* | succinate 30 °C 0 mM IPTG[a] | 2.505 | 2.841 |
| EL1548 | *p* $_{lac}$*-dnaA* | succinate 30 °C 0 mM IPTG[b] | 1.227 | 1.229 |
| EL558 | *p* $_{J23106}$*-dnaA* | succinate 30 °C | 0.973 | 0.997 |
| EL862 | *p* $_{J23100}$*-dnaA* | succinate 30 °C | 2.085 | 2.001 |
| EL864 | *p* $_{J23106}$*-dnaA ΔDARS1* | succinate 30 °C | 1.117 | 1.067 |
| EL868 | *p* $_{J23101b}$*-dnaA* | succinate 30 °C | 2.775 | 2.544 |
| EL2470 | *ΔDARS1* | succinate 30 °C | 1.070 | 1.060 |

[a] 90 min after switch from 1 mM IPTG.

[b] 180 min after switch from 1 mM IPTG.

[c] The same reference sample was split into two TMT-11 plex isobaric mass tagging reagents kits (samples A and B).

### Movie S1

***Movie S1.*** *Time-lapse of SeqA-Venus fluorescence and phase-contrast images before and after expression of DnaA was turned off. The frame rate was set to 15 frames per second.*
